# Structural basis of tRNA recognition by the m^3^C-RNA-methyltransferase METTL6 in complex with SerRS seryl-tRNA synthetase

**DOI:** 10.1101/2023.12.05.570192

**Authors:** Philipp Throll, Luciano G. Dolce, Palma Rico Lastres, Katharina Arnold, Laura Tengo, Shibom Basu, Stefanie Kaiser, Robert Schneider, Eva Kowalinski

## Abstract

Methylation of cytosine 32 in the anticodon loop of tRNAs to 3-methylcytosine (m^3^C) is crucial for cellular translation fidelity ^1^. Misregulation of the RNA methyltransferases setting this modification can cause aggressive cancers and metabolic disturbances ^2,3^. However, our understanding of the substrate selection and catalysis mode of the m^3^C RNA methyltransferases is currently still lacking. Here, we report the cryo-electron microscopy structure of the m^3^C tRNA methyltransferase METTL6 in complex with seryl-tRNA synthetase (SerRS) and their common substrate tRNA^Ser^. Through the complex structure, we identify the tRNA binding domain of METTL6. We show that SerRS acts as the tRNA^Ser^ substrate selection factor for METTL6. We reveal how METTL6 and SerRS jointly coordinate the long variable arm of tRNA^Ser^ in their interface. We demonstrate that SerRS augments the methylation activity of METTL6 and that direct contacts between METTL6 and SerRS are necessary for efficient tRNA^Ser^ methylation. Finally, based on the structure of METTL6 in complex with SerRS and tRNA^Ser^, we postulate a universal tRNA binding mode for m^3^C RNA methyltransferases including METTL2 and METTL8, suggesting that these mammalian paralogues use similar ways to engage their respective tRNA substrates and co-factors.

## Introduction

tRNAs play a central role in connecting mRNA codons to their corresponding amino acids and high-fidelity tRNA biogenesis is essential for accurate translation. Numerous nucleotide modifications ensure their correct folding and stability ^4^, protect them from nuclease cleavage ^5^, and ensure their correct subcellular localization ^6^. Moreover, nucleobase modifications in the tRNA anticodon loop expand the coding capacity of tRNAs, assure accurate decoding, and enhance translational fidelity and efficiency ^7–10^. 3-methylcytosine (m^3^C) present on base C_32_ in the anticodon loop of the majority of serine, threonine, and a subset of arginine isoacceptor tRNAs (tRNA^Ser^, tRNA^Thr,^ and tRNA^Arg^) is one of these nucleobase modifications ^1,11,12^. In the absence of m^3^C_32_, translation efficiency and fidelity are impaired ^3,13–15^.

In *Saccharomyces cerevisiae*, a single S-adenosyl-methionine (SAM)-dependent methyltransferase TRM140 (formerly ABP140) is responsible for m^3^C_32_ on tRNA^Thr^ and tRNA^Ser^^13,16^, while in the fission yeast *Schizosaccharomyces pombe* two distinct enzymes TRM140 and TRM141 target either threonine or serine isoacceptors, respectively ^17,18^. In mammals, three non-essential methyltransferases catalyze m^3^C on specific sets of tRNAs: METTL2A in cytosolic tRNA^Thr^ and tRNA^Arg^, METTL6 in cytosolic tRNA^Ser^ and METTL8 in mitochondrial (mt)- tRNA^Thr^ and mt-tRNA^Ser^ ^3,19–23^. These m^3^C RNA methyltransferases rely on co-factors to recognize and methylate their substrates or require pre-existing tRNA modifications for co-factor-independent substrate recognition. The molecular mechanisms of their substrate selection remain unclear.

In this study, we focused on METTL6, which is expressed in all tissues of the human body (https://www.proteinatlas.org/ENSG00000206562-METTL6). The enzyme has been found overexpressed in patients with highly proliferative luminal breast tumours and it is linked to poor prognosis in hepatocarcinoma ^2,3,24–28^. METTL6-deficient mice display lower metabolic rates and reduced energy turnover, and develop diverticular diseases ^3,29,30^. The methylation activity of METTL6 is specific for C_32_ of tRNA^Ser^. While there are reports of *in vitro* activity of isolated METTL6 ^3,31^, other studies suggested that METTL6 activity depends on seryl-tRNA synthetase (SerRS). Nevertheless, clear evidence for a direct interaction between both proteins is lacking, since their association was sensitive to RNase treatment ^19,20^. Recent crystal structures of METTL6 show how the enzyme interacts with the co-substrate SAM and the reaction co-product S-adenosyl homocysteine (SAH), but allowed only speculations about how METTL6 binds tRNA or interacts with SerRS ^31,32^.

Here, we report the cryo-electron microscopy (cryo-EM) structure of METTL6 bound to tRNA^Ser^ and SerRS at 2.4 Å resolution. Our data shows how the long variable arm of the tRNA^Ser^ is recognised by SerRS and anchors the tRNA in the interface between SerRS and METTL6. In addition, we reveal that METTL6 coordinates the tRNA anticodon arm through a domain that is common to all m^3^C RNA methyltransferases and remodels the anticodon stem-loop into a configuration that flips out the target nucleotide C_32_ rendering the base accessible for catalysis.

## Results

### SerRS boosts the catalytic activity of METTL6

We aimed to resolve the ambiguity about METTL6 dependence on SerRS for m^3^C methylation. For this, we expressed full-length human METTL6 and SerRS individually in *E. coli* and purified both proteins to homogeneity (Figure 1a, extended figure 1a-c). In our experiments, METTL6 was active on *in vitro* transcribed tRNA^Ser^ but not on tRNA^Thr^, used as a control. After adding SerRS to the reaction, the methyltransferase activity of METTL6 increased by ∼1000- fold. ATP and serine, the substrates of the aminoacylation reaction, were not necessary for the stimulation of METTL6 activity by SerRS (Figure 1b, extended Figure 1d). We thus conclude that SeRS acts as an activation factor of METTL6 that strongly increases methylation activity.

**Figure 1:**
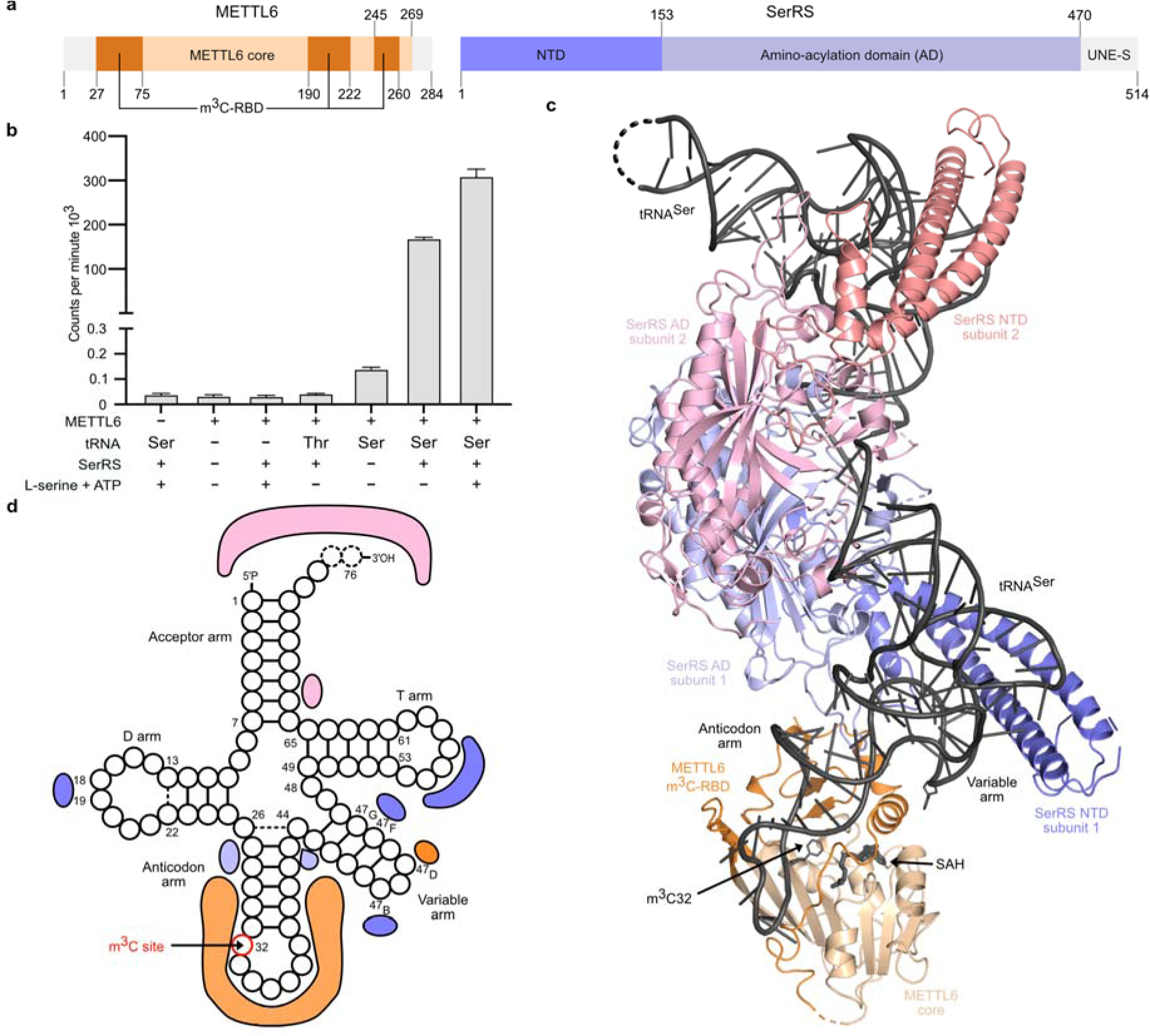
The structure of the SerRS-tRNA-METTL6 complex. a. Domain organization of human METTL6 and SerRS. The methyltransferase core of METTL6 is coloured in light orange and the RNA binding domain (m^3^C-RBD) in dark orange. The N-terminal domain (NTD) of SerRS is coloured in dark purple and the aminoacylation domain (AD) in light purple. Portions that are unresolved in the structure are coloured in grey. **b** *In vitro* methylation activity of METTL6 in the presence and absence of SerRS. Control experiments in Extended Figure 1d. Data is represented as the blank-subtracted mean and standard deviation of independent replicates (n=3). **c** The structural model of the 1:2:2 METTL6-SerRS-tRNA complex based on the cryo-EM reconstruction, protein domains are coloured as in panel a; in the second SerRS protomer the NTD is coloured in salmon and the AD in light pink.; tRNA and SAH are coloured in black. **d** Schematic depicting the protein contacts of METTL6 and SerRS to the commonly coordinated tRNA molecule. Colours as in panel **c**.

### METTL6 and SerRS form a tRNA-dependent complex

We aimed to assemble the METTL6-SerRS-tRNA complex to define how METTL6 recognizes tRNA^Ser^ and understand how METTL6 is activated by SerRS. Using size exclusion chromatography, we did not observe complex formation between purified METTL6 and SerRS. But the addition of tRNA^Ser^ led to the reconstitution of a trimeric complex (Extended Figure 1e). Despite attempts to crystallize the complex, we could only achieve two crystal forms of METTL6 bound to SAH (Extended Figure 1f and Extended Table 1) and the trimeric complex disintegrated during cryo-EM grid preparation. Therefore, we stabilised the assembly by fusing METTL6 to the flexible C-terminus of SerRS and co-expressed both proteins as a single polypeptide in insect cells. Under low-stringency purification conditions, the SerRS-METTL6 fusion construct was co-purified with cellular tRNA and subjected to cryo-EM structure solution (Extended Figure 2a).

**Figure 2:**
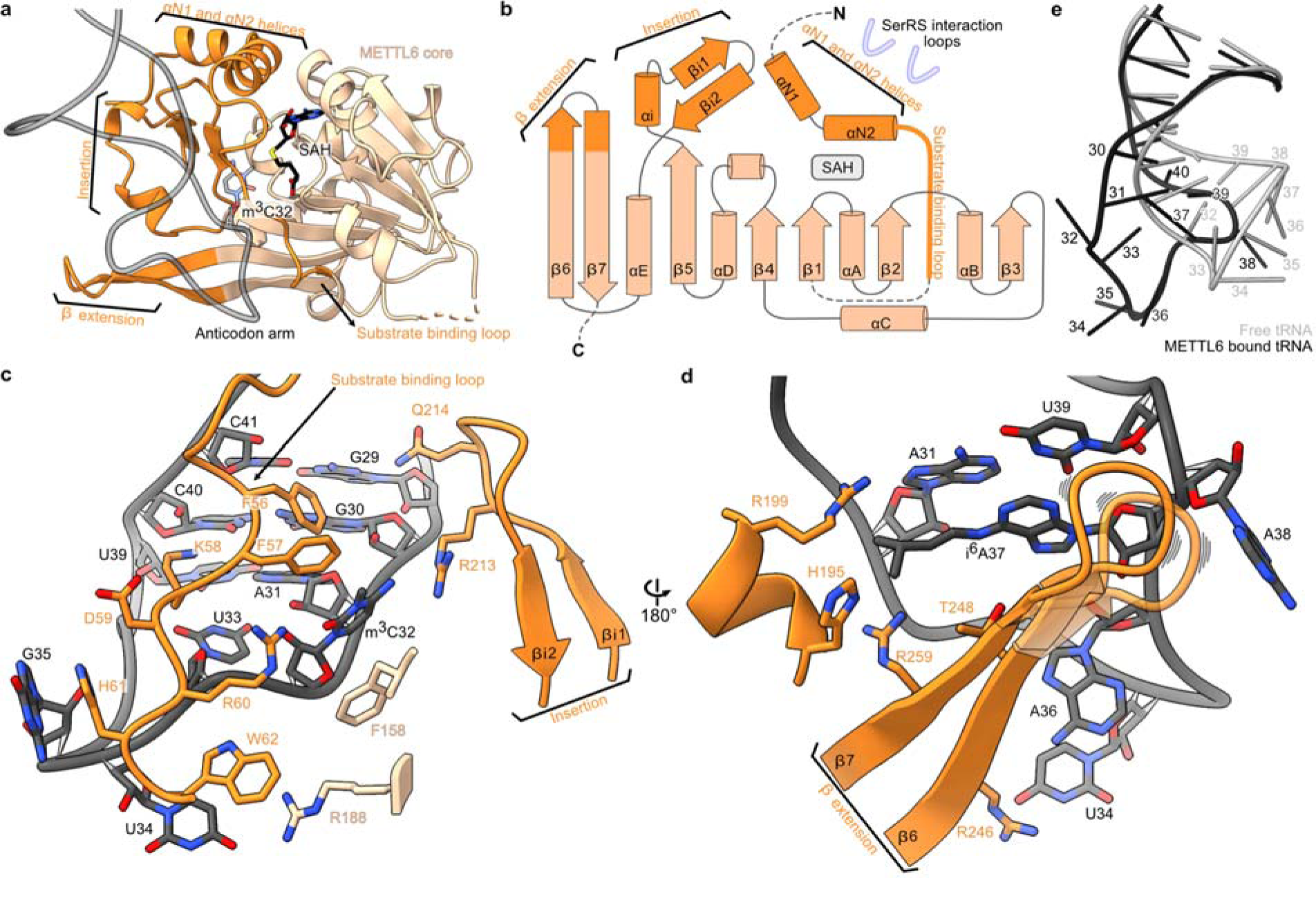
The m^3^C-specific RNA binding domain (m^3^C-RBD) of METTL6. a. Cartoon model of METTL6 from the complex structure with the methyltransferase core in light orange and the m^3^C-RBD in dark orange. The phosphodiester backbone of the anticodon arm of the tRNA in grey; C_32_ and SAH are represented as stick models. **b** Rossmann fold scheme of METTL6. The three segments of m^3^C, the SerRS interacting region and SAH binding site are indicated. **c** The substrate binding loop and β-turn part of the insertion contribute to binding of the anticodon loop of the tRNA. **d** The beta-elongation and alpha-helical part of the insertion embrace the anticodon loop from the opposite side and coordinate the i^6^A_37_ RNA modification. **e** Superposition of free tRNA (PDB:1ehz, gray) and the tRNA in the SerRS-METTL6 complex (black, rmsd = 1.124 Å). In all panels, the methyltransferase core is coloured in light orange and the m^3^C-RBD in dark orange; the RNA and SAH in shades of grey; stick models are coloured by heteroatom.

### Structural analysis of the METTL6-SerRS-tRNA^Ser^ complex

Processing of the cryo-EM micrographs and particle images of the METTL6-SerRS- tRNA^Ser^ complex revealed 3D classes for 1:2:1, 1:2:2 and 2:2:2 stoichiometries, partially because SerRS forms an obligatory dimer that can bind one or two tRNAs ^33–37^. Surprisingly, despite the fusion of both proteins, not all SerRS molecules in the reconstructions displayed density for associated METTL6. In the absence of protease treatment or cleavage sites, this suggests that the 60 amino acids long flexible linker connecting the proteins allowed free movement of METTL6 and that the fusion strategy tolerated an on-off equilibrium of the assembly (Extended Figure 2b). In the 3D class of the METTL6:SerRS:tRNA^Ser^ complex with a 2:2:2 stoichiometry (3.7 Å), the two copies of METTL6 are located approximately 100 Å apart from each other, each binding the proximal tRNA. This distance suggests that the minimal unit required for m^3^C-tRNA^Ser^ methylation is a METTL6 monomer acting on a single tRNA together with a SerRS dimer (Figure 1c and Extended Figure 2b). For model building, we therefore focused on the highest resolution class (2.4 Å), resolving a 1:2:2 METTL6: SerRS: tRNA complex (Extended Table 2).

In brief, at the core of the 1:2:2 METTL6:SerRS:tRNA complex, two molecules of tRNA^Ser^ bind symmetrically across the SerRS dimer without contact with each other. Each tRNA interacts with both SerRS protomers (Figure 1c, d). METTL6 is located at the extremity of the complex, binding its proximal tRNA anticodon stem and the variable arm that protrudes from the SerRS- tRNA^Ser^ complex. The tRNA^Ser^ long variable arm is coordinated in the interface between METTL6 and the SerRS N-terminal stalk. In the same interface, two protein loops of SerRS interact directly with METTL6.

### Insights into tRNA^Ser^ recognition by human SerRS

SerRS selects serine isoacceptors by binding their characteristic long variable arm with a helix of the N-terminal domain (NTD) stalk in the cleft between the T arm and the long variable arm (Figure 1c, d) ^33–39^. In contrast to previous human SerRS-tRNA^Sec^ complex structures ^37^, our METTL-SerRS-tRNA^Ser^ structure revealed a state in which the tRNA is fully engaged with the stalk of SerRS, resembling rather the tRNA-bound crystal structure of the *T. thermophilus* (Extended Figure 3a) ^33^. Therefore, our structure reveals unprecedented details of how human SerRS binds the variable arm of tRNA^Ser^ via nucleotide U_47B_ and discloses how the tRNA- geometry-defining 19:56 base pair is recognised (Extended figure 3b, c). The cryo-EM map does not resolve the full 3’-end of the acceptor arm, indicating that it is not engaged in the aminoacylation active site (Extended Figure 3d). Therefore, in addition to the insights gained into the METTL6-tRNA^Ser^-SerRS interaction, the complex structure reveals novel details of the binding of eukaryotic SerRS to tRNA^Ser^.

**Figure 3:**
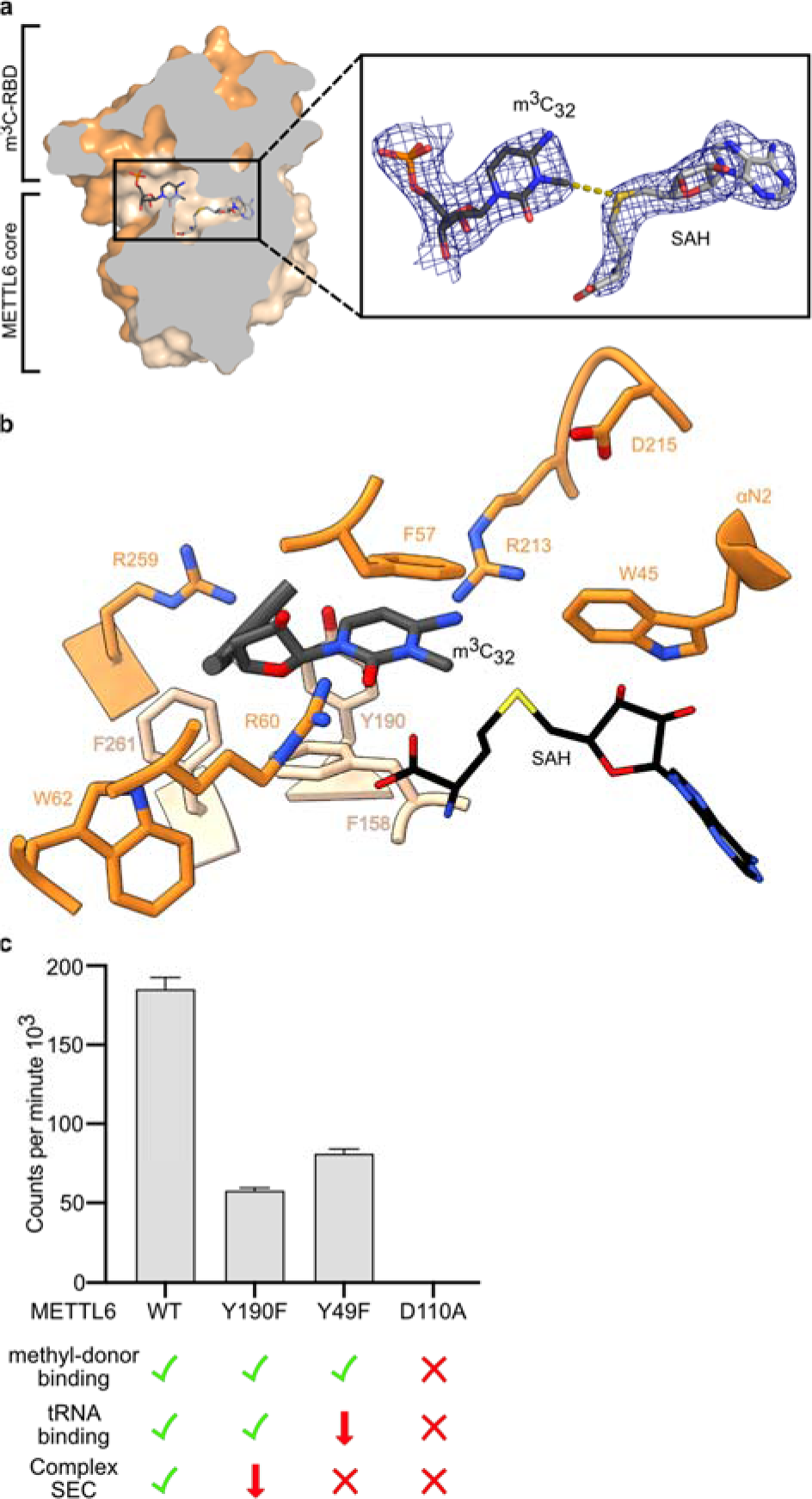
The METTL6 active site. a. Surface slice representation of METTL6 from the cryo- EM structure with the m^3^C_32_ and SAH in the active site, which lies in the interface between methyltransferase core and m^3^C-RBD. The zoomed-in area shows the EM map around m^3^C_32_ and SAH. **b** m^3^C_32_ coordination in m^3^C-RBD adjacent to SAH. **c** Methyltransferase activity of single point-mutants in the active site with a qualitative indication of control experiments for complex formation assessed via SEC (Extended Figure 7a) SAH binding by Thermofluor (Extended Figure 7c) and tRNA binding by FP (Extended Figure 7d). Methyltransferase control experiments in Extended Figure 7b. Data is represented as the blank-subtracted mean and standard deviation of independent replicates (n=3).

### The m^3^C-specific tRNA binding domain of METTL6 is conserved to the m^3^C RNA methyltransferase family

METTL6 shares the Rossmann-like fold of class I methyltransferases ^40,41^. The family of m^3^C-specific RNA methyltransferases (Interpro: IPR026113) have divergent N-terminal sequences and a conserved insertion within the Rossman fold, both of undetermined function (Extended Figure 4). Our structure of the METTL6-SerRS-tRNA^Ser^ complex pinpoints that three m^3^C methyltransferase-specific elements form a tripartite domain comprising the N-terminal region, the internal insertion, and the extended hairpin of the central ß-sheet. All elements of this “m^3^C-methyltransferase-specific RNA binding domain” (m^3^C-RBD) contribute to RNA binding. (Figure 2a, b).

**Figure 4:**
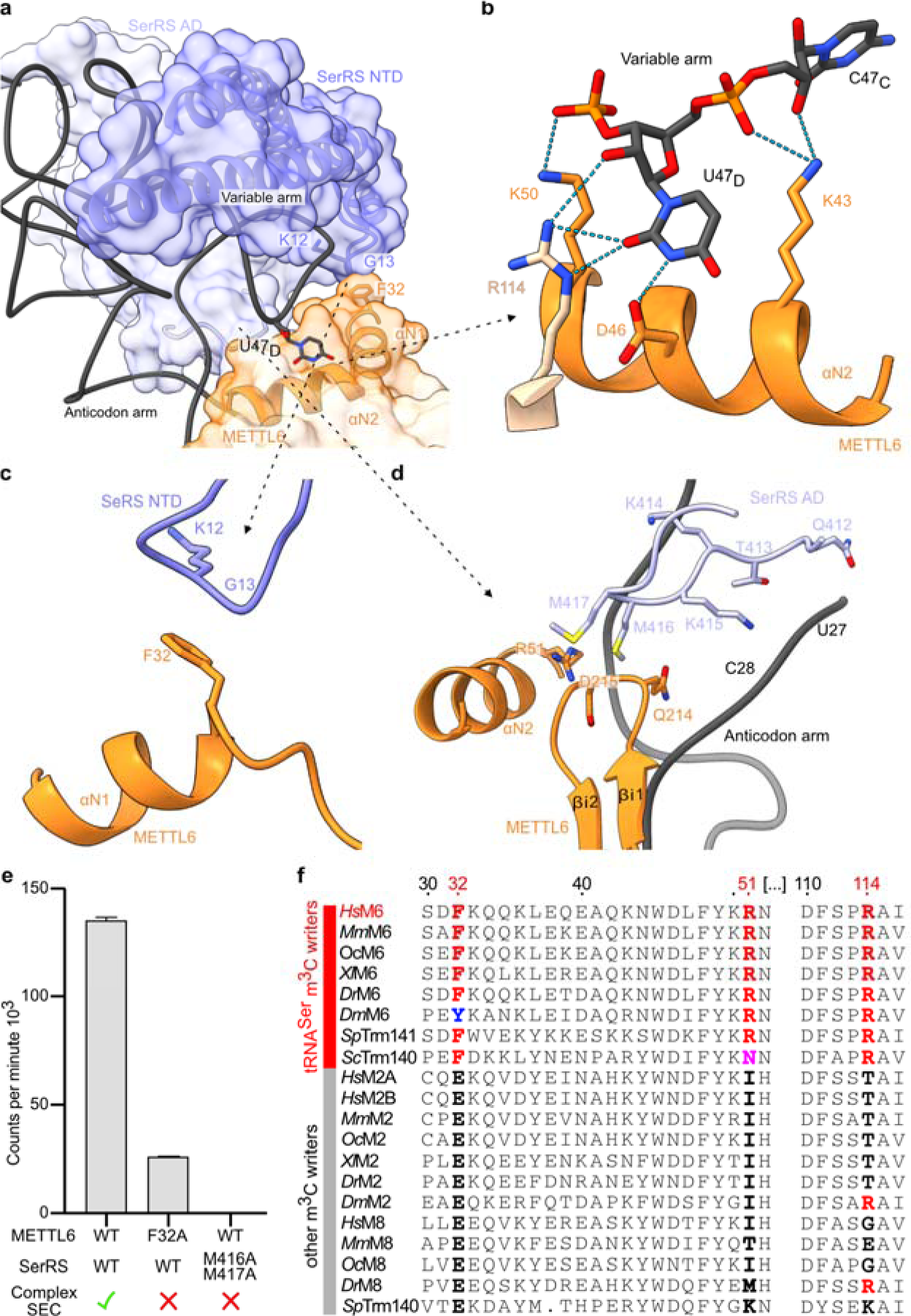
The METTL6-SerRS-tRNA variable arm interface. a. Cartoon and surface representation of the METTL6-SerRS-tRNA variable arm interface. SerRS NTD and αN1 and αN2 helices of METTL6 as cartoon representation with the F32 contact region to SerRS NTD loop indicated. U47D as stick representation. **b** Variable arm contacts to METTL6, cartoon and stick representation, dotted lines represent hydrogen bonds. **c** Interaction of METTL6 F32 in with the backbone of a SerRS NTD loop. **d** Interaction of METTL6 m^3^C-RBD with SerRS AD interface loop. Cartoon and stick representation. Colour scheme as in previous figures. **e** *In vitro* methyltransferase assay of interface mutants of METTL6 and SerRS with a qualitative indication of complex formation assessed via SEC (Extended figure 8e). Methyltransferase assay control experiments in Extended Figure 8f. Data is represented as the blank-subtracted mean and standard deviation of independent replicates (n=3). **f** Multiple sequence alignments of the variable arm interaction region of METTL6 with other m^3^C methyltransferases. Residues exclusively conserved in tRNA^Ser^-specific family members are highlighted in red.

In the N-terminal element of the m^3^C-RBD, two helices form the interface with SerRS in the complex. While the sequence of the first helix is not conserved across the m^3^C methyltransferase family members, the second helix (αN2) carries the conserved WDLFYK motif that we identify to play a central and multivalent role in coordinating the interactions with SerRS, SAH, and the variable arm of the tRNA. These helices are followed by an unstructured linker region carrying the strongly conserved FFKDRHW motif, which we termed the “substrate binding loop”. This motif conveys a negative charge to the binding cleft that accommodates the tRNA anticodon stem loop. The substrate binding loop is central in coordinating the target base C_32_ and contributes to the coordination of SAH (Figure 2c). The second element of the m^3^C-RBD is an insertion embedded between β5 and αE of the methyltransferase core comprising a short helical turn and a β-hairpin that both bind to the anticodon stem (Figure 2c, d). The third element of the m^3^C-RBD consists of a hairpin structure formed by the unusually elongated β-strands β6 and β7, which we termed β-extension. This hairpin region carries mostly positively charged residues, and the cryo-EM map indicates some degree of flexibility of this motif, possibly to enable the accommodation of diverse sequences in the variable anticodons of serine tRNAs (Figure 2d).

Surprisingly, the topological arrangement of the m^3^C-RBD of METTL6 did not resemble any other substrate-binding domains in characterised RNA or DNA methyltransferases. Among similarity search results, we identified the methyltransferases METTL11A and METTL11B that methylate the N-terminal α-amino group of proteins as relatives with a similar topology (Extended Figure 5a,b) ^41–46^ . Overall, our METTL6-SerRS-tRNA^Ser^ complex structure identifies the m^3^C- methyltransferase-specific RNA binding domain, which differs from so-far characterised DNA or RNA methyltransferases.

### tRNA^Ser^ binding induces structural rearrangement in the m^3^C-RBD

To determine whether tRNA binding to METTL6 induces conformational changes in the enzyme, we compared our structural model to the crystal structures of METTL6 bound to either SAM or SAH, but without an RNA ligand ^31,32^. The methyltransferase core of all models aligns well (Extended Table 3). Deviations occur exclusively in the m^3^C-RBD, in particular the divergent N-terminal region, the substrate binding loop, and the ß-extension. Most interestingly, all RNA-free structures display disordered substrate binding loops, which seems to adopt its shape only in contact with the tRNA, and a conserved helix following the substrate binding loop is rearranged. Our data suggest some degree of flexibility in the region of the ß-extension hairpin, which is completely unresolved in the crystal structures, suggesting its plasticity for tRNA^Ser^ accommodation (Extended Figure 5c). Thus, while the core of METTL6 is rigid, parts of the m^3^C- RBD seem more ductile to accommodate the tRNA.

### The METTL6-bound tRNA anticodon arm is remodelled for exposure of C_32_

Analysis of the cryo-EM map in the tRNA regions suggested that our sample contains a mixture of different serine isoacceptors. Accordingly, we built a tRNA^Ser^ consensus model based on the cryo-EM map density, tRNA^Ser^ sequence conservation, and the conservation of base modifications (Extended Figure 6a-d) ^11^. Comparing the METTL6-bound tRNA^Ser^ to the METTL6-unbound tRNA^Ser^ copy of our structure and also to free tRNA, we observe extensive remodelling of the anticodon arm and its global bending towards METTL6 (Figure 2e). In free tRNA, C_32_ is generally engaged in a stable non-canonical base pair with A_38 47_. Strikingly, in the complex structure, this C_32_:A_38_ base pair is disrupted and the bases are flipped to the outside of the anticodon loop. Such a flip-out mechanism as a prerequisite for base modification is a common feature in RNA-modifying enzymes such as ADAT and ADAR deaminases ^48–50^. Importantly, the extensive anticodon loop rearrangement brings C_32_ into the proximity of SAM bound in the METTL6 methyltransferase core and makes the base accessible for methyl transfer (Figure 2e).

### Coordination of the A_37_ modification

The cryo-EM map revealed a modification of adenosine A_37_ located in the anticodon loop.

Mass spectrometry identified predominantly i^6^A in the co-purified tRNA^Ser^ (Figure 2d and Extended Figure 6b,c). In the complex structure, the i^6^A_37_ residue base stacks with the last base pair of the anticodon stem (A_31_-U_39_); the modified isopentane moiety is coordinated by histidine H195, threonine T248, and two conserved arginine residues (R199 and R259), all located in the m^3^C-RBD (Figure 2d). The pre-modification of adenosine A_37_ to N6-isopentenyladenosine (i^6^A_37_) or N6-threonylcarbamoyladenosine (t^6^A_37_) has been described as a prerequisite for m^3^C_32_- modification in threonine tRNAs by METTL2A and METTL8 ^18,20–22^. Although it was not a requirement for METTL6 activity ^3,20^, the tight coordination of this modification in our complex structure suggests that i^6^A_37_ might assist in stabilizing the remodelled tRNA^Ser^ configuration. The two isopentane-coordinating arginine residues are conserved throughout the m^3^C methyltransferase family, suggesting that the METTL6 paralogues would coordinate i^6^A_37_ or t^6^A_37_ in a similar fashion (Extended Figure 4).

### m^3^C_32_ is coordinated through the methyltransferase core and m^3^C-RBD

The cryo-EM map density suggested the presence of SAH in our structure. In addition, mass spectrometry of the bound tRNA confirmed the presence of m^3^C-modified tRNA in our sample (Extended Figure 6b). In the structural model, the modified N3 atom of C_32_ lies at a 4 Å distance from the sulfur atom of SAH. These findings indicate that the structure captured a post-catalytic state with both products present (Figure 3a). The m^3^C_32_ nucleotide is tightly coordinated in the interface of the methyltransferase core and the m^3^C-RBD. Phenylalanines F158 and F57 stack the base via π-interactions, and arginine R60 forms a hydrogen bond to the carbonyl of the cytosine; arginine R213 also coordinates the base, and several other residues lie in distances that would allow van-der-Waals interactions to C_32_ (Figure 3b).

To probe the METTL6 active site experimentally, we tested single-point mutants for enzymatic activity, methyl-donor and tRNA^Ser^_UGA_ binding, and complex formation (Figure 3c, Extended Figure 7a-d). For methyl-donor binding, we used thermal stability assays in the presence of SAH, since this ligand resulted in the largest stability increase (Extended Figure 7b). To probe the tRNA- binding of METTL6 alone we conducted fluorescence polarization experiments in the absence of SerRS (Extended Figure 7c). The mutation of aspartate D110 to alanine (D110A), an ultra- conserved and indispensable SAM-coordination residue in this class of methyltransferases ^41^, resulted in the loss of SAH binding; it had impaired tRNA^Ser^_UGA_ affinity and was not active. The Y49F mutation, a tyrosine with a putative role in SAM binding, could still coordinate the methyl- donor, and showed weakened tRNA^Ser^ binding and attenuated activity. The mutation of tyrosine Y190 to alanine (Y190A), in proximity of m^3^C_32_, could still bind tRNA^Ser^_UGA_ but its catalytic activity was attenuated. All mutants were impaired in complex formation on size exclusion chromatography (Extended Figure 7d). As a side note, mutants of aspartate D189 to alanine or asparagine (D189A or D189N) were not soluble, suggesting a structural scaffolding role of this residue. We conclude that single-point mutations in the active site impair the interactions of METTL6 with tRNA^Ser^, the methyl-donor and SerRS, leading to attenuated methylation activity.

### Contacts between SerRS and METTL6 are indispensable for m^3^C_32_ modification of tRNA^Ser^

It was previously shown that METTL6 activity relies on the presence of the long variable arm of tRNA^Ser^ ^20^, but it was unclear how METTL6 recognizes this arm. The METTL6-SerRS- tRNA^Ser^ complex structure reveals that the variable arm of tRNA^Ser^ is embedded in the interface between METTL6 and SerRS (Figure 4a). METTL6 coordinates the strictly conserved variable arm residue U_47D_, through a hydrogen bonding network involving conserved residues of the m^3^C- RBD, such as aspartate D46 and lysine K50; arginine R114, in the core of METTL6, also contacts the variable arm (Figure 4b, Extended Figure 8a). Nevertheless, when we assessed tRNA binding in fluorescence polarization experiments, METTL6 alone could not distinguish tRNA^Ser^ from other tRNA species (Extended Figure 8b).

We hypothesised that rather SerRS selects the substrate and METTL6 recognizes a SerRS- tRNA^Ser^ complex. But remarkably only two loops of SerRS interact directly with METTL6 (Figure 4a). In the first contact point, the main chain of an NTD loop of SerRS contacts the side chain of phenylalanine F32 in the m^3^C-RBD (Figure 4c and Extended Figure 3a and 8c). The second SerRS interface loop, which is part of the aminoacylation domain, forms an intricate interaction network with several elements of the METTL6 m^3^C-RBD and also contacts the upper part of the tRNA anticodon stem (Figure 4d, Extended Figure 3a, 8d). Mutation of the METTL6 phenylalanine F32 to alanine (F32A) prevented complex formation in size exclusion chromatography and attenuated methylation activity (Figure 4e and Extended Figure 8e, f). The double mutant of the SerRS methionines to alanines (MM416/417AA) disrupted the complex and suppressed methylation activity (Figure 4e and Extended Data 8e, f). It is important to note that the METTL6 SerRS- interface residues phenylalanine F32 and arginine R51 are exclusive to m^3^C-methyltransferases that target cytosolic tRNA^Ser^ and that the SerRS contact loops are conserved as well (Figure 4f, Extended Figure 8g, h). Thus SerRS is the substrate recognition factor for METTL and the conserved direct contacts between METTL6 and SerRS are indispensable for the coordinated binding of the variable arm of tRNA^Ser^.

## Discussion

In this work, we present the cryo-EM structure of METTL6 in complex with SerRS bound to the co-products of the methyl-transfer, the m^3^C_32_-modified tRNA^Ser^ and SAH. We identify the tripartite tRNA-binding domain of METTL6 and show that SerRS is necessary as a tRNA- selection factor that augments METTL6 activity.

Functional key residues in the m^3^C-RBD of METTL6 are conserved to METTL2 and METTL8, such as the SAM-binding motifs, the substrate binding loop, the C_32_ coordinating residues and residues that coordinate the i^6^A_37_ or t^6^A_37_ modification (Extended Figure 4). These similarities suggest resembling RNA binding modes for all paralogues in the m^3^C RNA methyltransferase family. Indeed AlphaFold2 models for the baker’s and fission yeast homologs of METTL6 and the other human m^3^C_32_ methyltransferases METTL8 and METTL2A (https://alphafold.ebi.ac.uk/) are rather similar to the tRNA-bound structure of METTL6 (Extended Table 3). However, they are distinct in their divergent N-terminal sequences. Our data show that this N-terminal region of METTL6 has a dual role in mediating co-factor-specific interactions with SerRS and in providing substrate-specific interactions to the long variable arm of tRNA^Ser^. Analogously, METTL2 and METTL8 might carry specificity signals in their N-termini, e.g. for recruitment of mitochondrial SerRS2 to METTL8 for mt-tRNA^Ser^ selection, or for recruitment of DALRD3 to METTL2B for tRNA^Arg^ modification. Notably, DALRD3 is predicted to carry an anticodon binding domain similar to class Ia aminoacyl-tRNA synthetases. Our discovery of the tripartite configuration of the m^3^C-RBD shows that the three elements of the METTL6 m^3^C-RBD act collectively for SerRS binding and tRNA^Ser^ recognition. This observation clarifies why previous attempts to swap N-termini between METTL6 and METTL2A in chimeric enzymes did not change their substrate specificities ^20^.

Although the genes for METTL2, METTL6 and METTL8 are non-essential, all three enzymes have been linked to disease. METTL6 acts as an oncogene in luminal breast tumours and hepatocarcinoma ^2,3,24–28^, and METTL2A also seems to play a role in breast cancer ^51^. Similarly, the mitochondrial m^3^C methyltransferase METTL8 has been linked to severe pancreatic adenocarcinoma and colon cancer and is involved in neurogenesis ^22,52–54^. Our data reveal how m^3^C methyltransferases interact with RNA. Hence we establish a foundation for future detailed investigations on the enzymes of this family but also reveal promising prospects for potential therapeutic applications.

## Methods

### Cloning, protein expression and purification in *E. coli*

Protein coding DNA sequences for METTL6 (UniProt-ID: Q8TCB7) with a 3C-protease cleavable N-terminal 6His-GST tag and SerRS (UniProt-ID: P49591) with a C-terminal 6His-tag were cloned into a pEC vector. Point mutations of the proteins were produced by site-directed mutagenesis using single or double-mismatch primers.

All proteins were expressed in *E. coli* of the Rosetta II (DE3) strain and cultured in terrific broth medium. Expression was induced with 0.1 mM isopropyl-1-thio-β-d-galactopyranoside (IPTG) at 18°C overnight. Cell pellets were frozen and kept at -80°C before purification.

Wild type or mutant METTL6 was purified by resuspending and sonicating the cell pellet in lysis buffer containing 20 mM TRIS-HCl pH 7.5, 250 mM NaCl, 20 mM Imidazole, 0.05% (v/v) 2-mercaptoethanol, 1 mM PMSF, 1 μg/mL DNAseI, 0.5 μg/mL RNAse and 50 μg/mL lysozyme. The lysate was cleared by centrifugation with 20,000 x g followed by filtration with a 5 μM filter and loaded on a HisTrap HP column pre-equilibrated with lysis buffer (GE Healthcare). The column was washed first with lysis buffer, then with high-salt buffer (20 mM TRIS-HCl pH 7.5, 250 mM NaCl, 1 M KCl, 50 mM Imidazole) and low-salt buffer (20 mM TRIS-HCl pH 7.5, 100 mM NaCl, 100 mM Imidazole) before eluting with elution buffer (20 mM TRIS-HCl pH 7.5, 100 mM NaCl, 300 mM Imidazole). The protein was further purified by loading it on a HeparinTrap (GE Healthcare) and eluted with a gradient from 100 mM to 1 M NaCl. Protein-containing fractions were pooled, recombinant His-tagged 3C-protease (EMBL PEPcore) was added, and the protein was dialyzed overnight against 20 mM HEPES-KOH pH 7.5, 100 mM NaCl, 2 mM MgCl_2_, 20 mM Imidazole, 0.05% (v/v) 2-mercaptoethanol. The cleaved tag and 3C- protease were removed by running the dialysate over another 5 mL HisTrap column and collecting the unbound protein. The protein was concentrated using centrifugal spin concentrators (Millipore) with an MWCO of 10 kDa. As the final purification step size exclusion chromatography was performed on a Superdex 75 16/600 column (GE Healthcare) with 20 mM HEPES-KOH pH 7.5, 100 mM NaCl, 2 mM MgCl_2_, 1 mM DTT. Fractions with pure protein were pooled and concentrated. All protein purification steps were carried out at 4°C.

SerRS was purified by resuspending the cell pellet in lysis buffer (20 mM TRIS-HCl pH 7.5, 250 mM NaCl, 20 mM Imidazole, 0.05% (v/v) 2-mercaptoethanol, 1 mM PMSF, 1 μg/mL DNAseI, 0.5 μg/mL RNAse and 50 μg/mL lysozyme) and lysed by microfluidisation with a pressure of 275 kPa. The lysate was cleared by centrifugation with 20,000 x g at followed by filtration with a 5 μM filter before loading on a HisTrap HP column (GE Healthcare). The column was washed first with lysis buffer, then with high-salt buffer (20 mM TRIS-HCl pH 7.5, 250 mM NaCl, 1 M KCl, 50 mM Imidazole) and low-salt buffer (20 mM TRIS-HCl pH 7.5, 100 mM NaCl, 50 mM Imidazole) before eluting with elution buffer (20 mM TRIS-HCl pH 7.5, 100 mM NaCl, 300 mM Imidazole). The eluted protein was then applied to a 5 mL Q-trap (GE Healthcare) and eluted with a gradient from 100 mM to 1 M NaCl. Pure protein-containing fractions were pooled and concentrated using centrifugal spin concentrators with an MWCO of 30 kDa (Millipore). Finally, SEC was performed on a Superdex 200 10/300 column with 20 mM HEPES- KOH pH 7.5, 100 mM NaCl, 2 mM MgCl_2_, 1 mM DTT. Fractions containing the pure protein were pooled and concentrated. The purified proteins were flash-frozen in liquid nitrogen and stored at -80°C for further experiments.

### Cryo-EM sample preparation

The DNA sequences coding for SerRS and METTL6 were cloned into a psLIB vector by Gibson Assembly, yielding a fusion construct with the sequence for METTL6 fused directly to the C-terminus of SerRS with a C-terminal 3C-protease cleavable EGFP tag. The fusion protein was expressed in Hi5 cells. After 72h of protein expression at 25°C, the cells were collected by centrifugation, resuspended in lysis buffer (20 mM TRIS-HCl pH 7.5, 100mM NaCl, 2mM MgCl_2_, 0.05% (v/v) 2-mercaptoethanol, 1 mM PMSF) and lysed by sonication. The lysate was cleared by centrifugation with 20,000 x g at 4°C followed by filtration with a 5 µM filter. All subsequent purification steps were carried out at 4°C.

The lysate was incubated with 1 mL EGFP nanobody resin for 30 minutes. The resin was washed two times with a wash buffer containing 20 mM TRIS HCl pH 7.5, 100 mM NaCl and 2 mM MgCl2. The SerRS-METTL6 fusion construct was eluted from the resin by cleavage of the eGFP-tag using the 3C-protease cleavage site: 500 µL wash buffer with 0.08 mg/mL 3C-protease (EMBL PEPcore) was added to the resin and incubated for circa 2 hours. The cleaved protein was removed from the resin, filtered with a Spin-X centrifuge tube filter (Costar) with a pore size of 0.22 µM and concentrated using a centrifugal spin concentrator (Millipore) with an MWCO of 10 kDa. As the final purification step, the protein was purified by size exclusion chromatography on a Superdex 200 3.2/300 column equilibrated in 20 mM HEPES-KOH pH 7.5, 100 mM NaCl, 2 mM MgCl_2_, 0.5 mM TCEP. Fractions containing the fusion construct bound to tRNA were frozen with liquid nitrogen and stored at -80°C until grid preparation.

The sample was diluted to 0.2 mg/mL with a buffer containing 20 mM HEPES-KOH (pH 7.5), 50 mM NaCl, 2 mM MgCl_2_, 0.5 mM TCEP and 1 mM sinefungin. For electron microscopy grid preparation, UltrAuFoil grids with an R 1.2/1.3 300 mesh (EMS) were glow-discharged on both sides for 40 seconds with 30 mA at 0.45 bar using a Pelco EasyGlow glow-discharging device. 2 µL of the diluted sample were applied to each side on the grid, and the sample was vitrified in liquid ethane using a MARK IV Vitrobot (FEI) with the following settings: 100% humidity, 4°C, 3 seconds blot time and a blot force of 0.

### Cryo-EM data acquisition, processing, and model-building

Data was acquired on a Titan Krios transmission electron microscope (FEI) equipped with a 300 kV accelerating voltage field emission gun electron source, a Quantum energy filter (Gatan) and a Gatan K3 camera, at 130K magnification with a pixel size of 0.645 Å/pixel. The microscope was operated using the software SerialEM. Movies were acquired in counting mode with an electron dose of 63.27 e^-^/Å^2^ in 40 frames with a defocus range from -0.8 µm to -1.8 µm.

Motion correction and CTF estimation on the collected micrographs was performed using RELION 3.1 ^55^. Particles were picked with the BoxNet_20220403_191253 model in Warp ^56^, and extracted in RELION in a 550 pixel box. A total of 1,625,946 particles were used as input for data processing using cryoSPARC ^57^. After an initial round of 2D classification to remove bad particles and particles containing only tRNA, the remaining 1,282,076 particles were used for several rounds of *ab initio* 3D classifications, which resulted in different classes containing dimeric SerRS with one or two tRNA molecules and 0-2 copies of METTL6 bound (Extended Figure 2 and Extended Table 2). The particles of the 3D class with dimeric SerRS, two tRNAs and one molecule of METTL6 were subjected to several rounds of CTF refinement and Bayesian polishing in RELION 3.1. The refined particles were re-imported into cryoSPARC to perform a final non- uniform refinement ^58^. All reported resolutions were determined using the gold standard FSC method implemented in cryoSPARC, with a cutoff of 0.143.

The highest resolution model was built in coot using the available crystal structures of tRNA-bound human SerRS (PDB-ID: 4RQE ^37^) and of METTL6 (this study) as starting templates. The model was refined by several rounds of real-space refinement in Phenix (version 1.16) and manual refinement in Coot (version 0.8.9.2). The other models were build based on the highest resolution model, with minimal refinement.

### Protein crystallography

METTL6 was crystallised by vapour diffusion in the hanging drop format at 20°C at a concentration of 4 mg/mL supplanted with 2 mM SAH and 5 μM TCEP. 1 μL of protein solution was added to 1 μL of reservoir solution containing 0.1 M BisTRIS propane pH 6.5, 0.2 M sodium sulfate, and 20 % PEG3350. After two weeks crystals were soaked for two minutes in a cryoprotectant solution containing 2 mM SAH, 0.1 M BisTRIS propane pH 6.5, 0.2 M sodium sulfate, 10% PEG400 and 30% PEG 3350 and flash-frozen in liquid nitrogen.

A truncated construct of METTL6 comprised of residues 40 to 269 was crystallised at a concentration of 5 mg/mL supplanted with 1 mM SAH in the sitting drop format platform by adding 100 nL protein solution to 100 nL of a reservoir solution containing 0.1 M BisTRIS, 0.2 M ammonium sulfate and 25% PEG3350 at 20°C. The crystals were harvested using the automated Crystal Direct harvesting system of the HTX platform ^59^ and flash-frozen without cryoprotectant. X-ray diffraction data sets of full-length METTL6 were collected on the MASSIF-1 beamline at the ESRF with an x-ray energy of 12.65 keV (= 0.9801 Å). For solving the phase problem for METTL6 by single-wavelength anomalous dispersion (SAD) with the anomalous signal from sulfur, a total of 35 x-ray diffraction datasets with an x-ray energy of 6 keV (= 2.066 Å) were collected on beamline P13 at the PETRA-III storage ring at DESY. X-ray diffraction data sets of METTL6Δ were also obtained on this beamline with an x-ray energy of 12.3 keV (= 1.008 Å).

All diffraction images were processed using XDS ^60^. For solve the phase problem, five of the 6 keV datasets were clustered and merged using blend ^61^ and fed into the CRANK2 pipeline for SAD phasing ^61^ which resulted in a low resolution (4.42 Å) structure of METTL6. This low-resolution structure was used to obtain phases for the other data sets by molecular replacement using Phenix-phaser ^62^. For all models, several rounds of model building using COOT ^63^ and data refinement using PHENIX-REFINE ^64^ were performed. Ramachandran statistics for the METTL6 FL are 97.42% favoured, 2.58% allowed and 0% disallowed and for the METTL6 40-269) are 95.18% favoured, 4.82% allowed and 0% disallowed. Data collection and refinement statistics are shown in Extended Table 1.

### tRNA *in vitro* transcription

The DNA sequences coding for human tRNA^Ser^ and tRNA^Thr^ were cloned into a pUC19 vector between a T7 promoter and a BstN1 cleavage site. The vector was linearised using BstN1 and used as a template for runoff transcription with T7 polymerase. 1 mg of linearised DNA was *in vitro* transcribed in a reaction containing 40 mM TRIS pH 8.0, 30 mM MgCl_2_, 5 mM DTT, 1 mM spermidine, 0.01 % Triton X100, 4 mM of each NTP and 50 µg/mL recombinant T7 polymerase (EMBL PEPcore), which was incubated for 16 h at 37°C. The produced RNA was isopropanol precipitated, purified via Urea-PAGE, desalted using Pierce dextran desalting column (ThermoFischer) and concentrated using a centrifugal spin concentrator (Millipore) with a MWCO of 10 kDa. The tRNA was refolded by heating it for 2 minutes to 95°C before adding MgCl_2_ to a final concentration of 2 mM and quickly placing it on ice.

### Thermal Shift Assay

METTL6 was diluted to a final concentration of 20 μM in a buffer containing 20 mM HEPES-KOH (pH 7.5), 100 mM NaCl, 2 mM MgCl2, 1 mM DTT, 5X SYPRO Orange (Invitrogen) and 1 mM of cofactor (SAM, SAH or sinefungin). Fluorescence was measured in a real-time PCR machine (Stratagene Mx3005P) while subjected to a temperature gradient from 24.6°C to 95°C in steps of 1°C and 1 min.

### Fluorescence polarization assay

The *in vitro* transcribed tRNA^Ser^ and tRNA^Thr^ were labelled on the 3’ end with fluorescein. The DNA, and RNA oligomers (labelled on the 3’-end with 6-carboxyfluorescein) were purchased from biomers.net. All nucleic acids were refolded as described above. The labelled tRNAs with a concentration of 25 nM were incubated with METTL6 of different concentrations in a total volume of 20 µL for 15 minutes at room temperature in a 384-well flat black microplate (Greiner) in a buffer of 20 mM HEPES-KOH (pH 7.5), 50 mM NaCl, 2 mM MgCl_2_, 0.1 % TWEEN, 1 mM DTT, 0.5 U/µl RNasin and 0.5 mM Sinefungin. Fluorescence polarisation was measured at a temperature of 25°C with a ClarioStar plate reader (BMG labtech) with an excitation wavelength of 460 nm and an emission wavelength of 515 nm. Data analysis was performed using GraphPad Prism using the single site binding fit.

### Size-exclusion complex formation assay

METTL6, SerRS and tRNA^Ser^ (each at a concentration of 30 µM; METTL6 supplemented with 1 mM sinefungin to stabilize the complex) were incubated for 10 minutes on ice before subjecting them to analytical size exclusion chromatography using a Superdex 200 Increase 3.2/300 column (GE Healthcare) equilibrated in a buffer composed of 20 mM HEPES-KOH pH 7.5, 50 mM NaCl, 2 mM MgCl_2_, 0.5 mM TCEP. Fractions were analyzed via SDS-PAGE stained with coomassie blue and Urea-PAGE stained with methylene blue for protein and RNA content respectively.

### Methyltransferase assay

Methyltransferase assays were performed in 6 mM Hepes-KOH (pH 7.9), 0.4 mM EDTA, 10 mM DTT, 80 mM KCl, 1.5 mM MgCl_2_, RNasin (40 U/µl) (Promega), and 1.6% glycerol. 0.05 µM of the corresponding enzyme was used, and 0.4 µM of *in vitro* transcribed tRNA(ser) or tRNA(thr) or 5ug of total RNA extracted from Hela cells was added as a substrate. 4 µM of Serine (Sigma) was added where indicated. 1 µl of ^3^H-labelled SAM (1 mCi/ml; PerkinElmer) was added to the mixture and then incubated at 30°C for 1 h with gentle shaking. Another 1 µl of ^3^H-SAM was added and the assay continued at room temperature (22°C) overnight followed by column purification with the Zymo Research RNA miniprep kit. Tritium incorporation was analysed by liquid scintillation counting with a Triathler counter (Hidex) after adding the Filter-Count liquid scintillation counter cocktail (PerkinElmer). All data of every *in vitro* methyltransferase assay are shown as blank-substracted mean of three replicates, and three experimental replicates were carried out for each experiment.

### Sample preparation for LC-MS/MS and detection of modified nucleosides

The *Trichoplusia ni* tRNA that co-purified with the SerRS-METTL6 fusion construct was isolated via chloroform-phenol extraction and precipitated with ethanol and sodium acetate. Around 300 ng of the tRNA was digested in an aqueous digestion mix (30 μL) to single nucleosides by using 2 U alkaline phosphatase, 0.2 U phosphodiesterase I (VWR, Radnor, Pennsylvania, USA), and 2 U benzonase in Tris (pH 8, 5 mM) and MgCl_2_ (1 mM) containing buffer. Furthermore, 0.5 μg tetrahydrouridine (Merck, Darmstadt, Germany), 1 μM butylated hydroxytoluene, and 0.1 μg pentostatin were added. After incubation for 2 h at 37 °C, 20 μL of LC-MS buffer A (QQQ) was added to the mixture. A stable isotope labelled SILIS (gen², ^65^ was added to each replicate and calibration solution of synthetic standards before injection into the QQQ MS.

### LC-MS/MS of nucleosides

Modified nucleosides were identified and quantified using mass spectrometry. An Agilent 1290 Infinity II equipped with a diode-array detector (DAD) combined with an Agilent Technologies G6470A Triple Quad system and electrospray ionization (ESI-MS, Agilent Jetstream) was used.

Nucleosides were separated using a Synergi Fusion-RP column (Synergi® 2.5 μm Fusion-RP 100 Å, 150 × 2.0 mm, Phenomenex®, Torrance, CA, USA). LC buffer consisting of 5 mM NH_4_OAc pH 5.3 (buffer A) and pure acetonitrile (buffer B) were used as buffers. The gradient starts with 100% buffer A for 1 min, followed by an increase to 10% buffer B over a period of 4 min. Buffer B is then increased to 40% over 2 min and maintained for 1 min before switching back to 100% buffer A over a period of 0.5 min and re-equilibrating the column for 2.5 min. The total time is 11 min and the flow rate is 0.35 mL/min at a column temperature of 35 °C.

An ESI source was used for ionisation of the nucleosides (ESI-MS, Agilent Jetstream). The gas temperature (N2) was 230 °C with a flow rate of 6 L/min. Sheath gas temperature was 400 °C with a flow rate of 12 L/min. Capillary voltage was 2500 V, skimmer voltage was 15 V, nozzle voltage was 0 V, and nebuliser pressure was 40 Psi. The cell accelerator voltage was 5 V. For nucleoside modification screening, MS2Scan (*m/z* 250-500) was used. For quantification, a DMRM and positive ion mode was used (Extended data Table 4).

### Data analysis of nucleosides

For calibration, synthetic nucleosides were weighed and dissolved in water to a stock concentration of 1-10 mM. Calibration solutions ranged from 0.0125 pmol to 100 pmol for each canonical nucleoside and from 0.00625 pmol to 5 pmol for each modified nucleoside. The concentrations of Ψ and D ranged from 0.00625 pmol to 20 pmol. Analogous to the samples, 1 µL of SILIS (10×) was co-injected with each calibration. The calibration curve and the corresponding evaluation of the samples were performed using Agilent’s quantitative MassHunter software. All modification abundances were normalised to the amount of RNA injected using the sum of all canonicals.

## Data availability

The cryo-EM maps and their derived models for the deposited in the PDB: accession code 8P7B for the 1:2:2 METTL6:SerRS:tRNA stochiometry, 8P7C for the 2:2:2 complex and 8P7D for the 1:2:1 complex. EMDB entries in the same order are EMD-17528, EMD-17529 and EMD- 17530. Maps of SerRS-tRNA and tRNA 3D-classes were side products of the data processing and have been deposited under EMD-17531 and EMD-17532. All micrographs of the dataset are deposited at EMPIAR-XXXXX. The crystal structures of METTL6 are deposited in the PDB (8OWX and 8OWY). Source data are provided in addition to this paper.

### Acknowledgements

We acknowledge Martin Pelosse for support in using the Eukaryotic Expression Facility at EMBL Grenoble, and Sarah Schneider for support in using the EM Facility at EMBL Grenoble. This work benefited from access to the cryo-EM platform of the Structural and Computational Biology Unit at EMBL Heidelberg and we thank Félix Weis for his assistance with the data collection. We acknowledge the HTX Team for crystallization setups, automated crystal harvesting and thermostability assays. This work used the platforms of the Grenoble Instruct-ERIC Centre (ISBG; UAR 3518 CNRS-CEA-UGA-EMBL) within the Grenoble Partnership for Structural Biology (PSB), supported by FRISBI (ANR-10-INBS-0005-02) and GRAL, financed within the University Grenoble Alpes graduate school (Écoles Universitaires de Recherche) CBH-EUR-GS (ANR-17-EURE-0003). Work in the R.S. laboratory was supported by the DFG through SFB 1064 (Project-ID 213249687) and the Helmholtz Gesellschaft. Both R.S. and S.K. are supported through SFB 1309 (Project-ID 325871075). We thank Life Science Editors for editing services (www.lifescienceeditors.com). The authors thank the Kowalinski lab members for discussions and comments throughout the course of the project.

## Author contributions

EK conceptualised and led the study. PT and LT conducted protein purifications, RNA *in vitro* transcription, SEC experiments, FP, cryo-EM sample preparation. SB assisted in x-ray data collection, phasing and data processing. PRP, KA conducted activity assays in the lab of RS. SK conducted nucleoside modification mass spectrometry. PT and LGD collected and processed cryo- EM data. PT, LGD and EK built the models and interpreted the data. EK wrote the manuscript supported by LGD, RS and PT.

## Extended Data

**Extended Data Figure 1:**
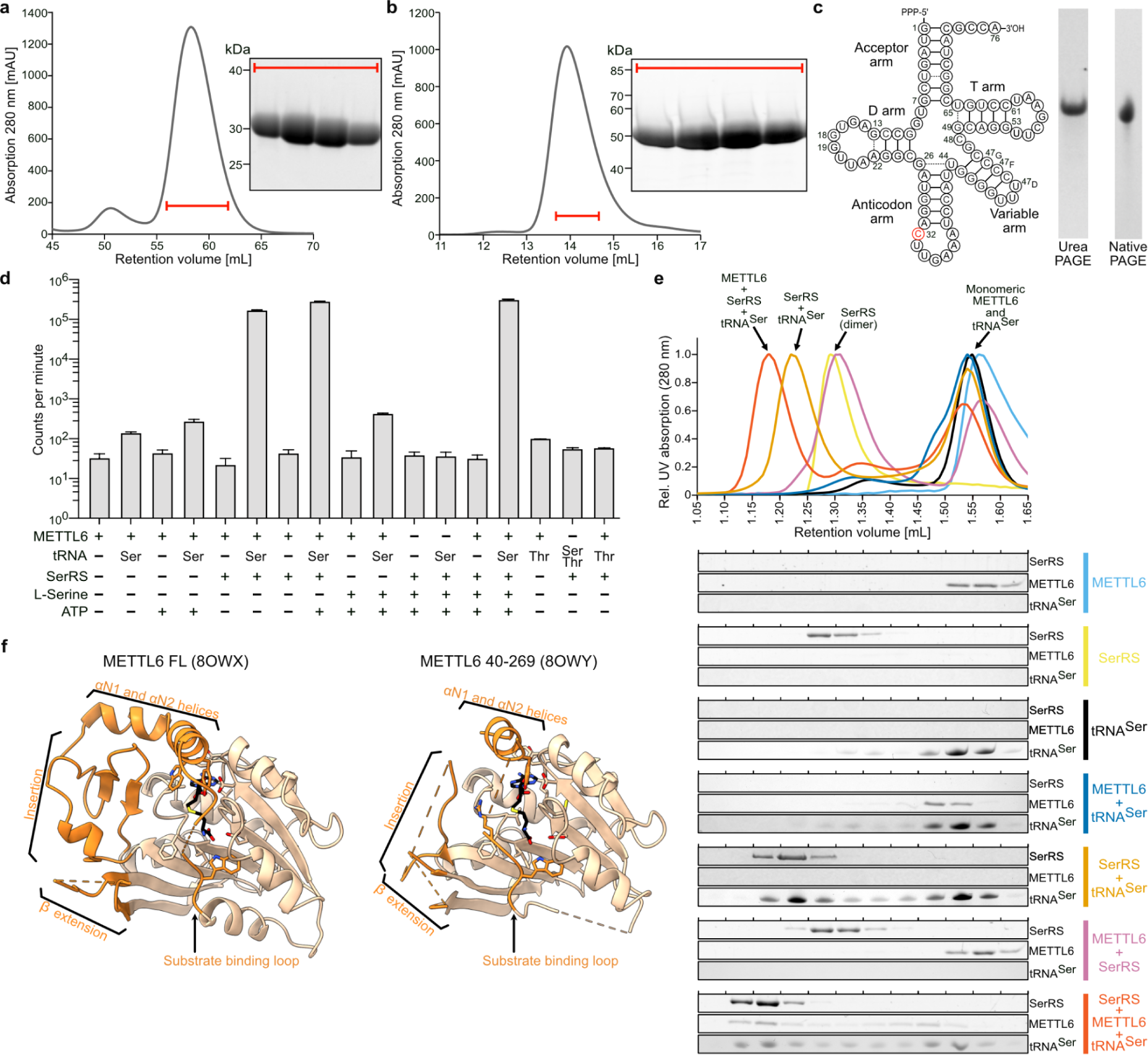
Protein/RNA purification and cryo-EM sample preparation **a** Size-exclusion chromatogram and SDS-PAGE analysis of the last step of the purification of full-length METTL6 (Column: Superdex 75 16/600). **b** Size-exclusion chromatogram and SDS-PAGE analysis of the last step of the purification of full-length SerRS (Column: Superdex 200 10/300). **c** Sequence (left) and quality control of *in vitro* transcribed tRNA^Ser^ on denaturing urea PAGE (left) and native PAGE (right). **d** *In vitro* methylation activity measurements of METTL6 in the presence of SerRS, tRNA^Ser^ and/or L-Serine and ATP. Data is represented as the blank-subtracted mean and standard deviation of independent replicates (n=3). **e** Complex formation of SerRS, tRNA^Ser^ and METTL6 on analytical size-exclusion chromatography (Column: Superdex 200 3.2/300). Fractions were analysed for protein and RNA contents using SDS-PAGE and urea PAGE. **f** Cartoon representation of crystal structures from this study: apo-METTL6 full-length (pdb:8OWX) and apo-METTL6 40-269 (pdb 8OWY) in complex with SAH.

**Extended Data Figure 2:**
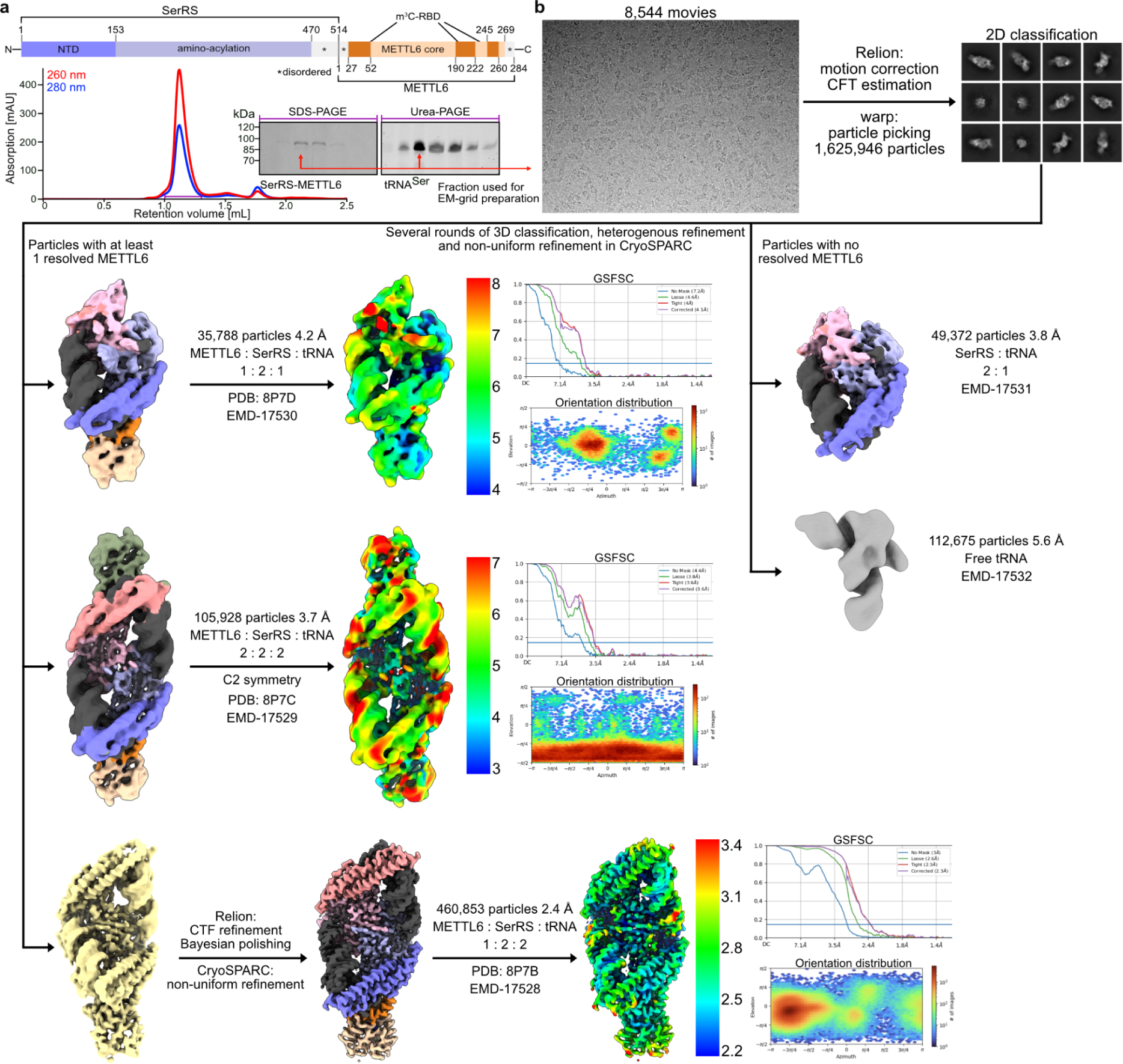
Cryo-EM data processing scheme. **a** Cryo-EM sample preparation with a schematic of the SerRS-METTL6 fusion polypeptide. Size-exclusion chromatogram and SDS-PAGE and urea PAGEs of a typical purification are depicted. The fraction marked with a red arrow was subjected to cryo-EM grid vitrification. **b** Cryo-EM data processing scheme. The final resolution estimates were calculated using the gold standard FSC method implemented in CryoSPARC. Colouring of the structure by components as in main Figure 1.

**Extended Data Figure 3:**
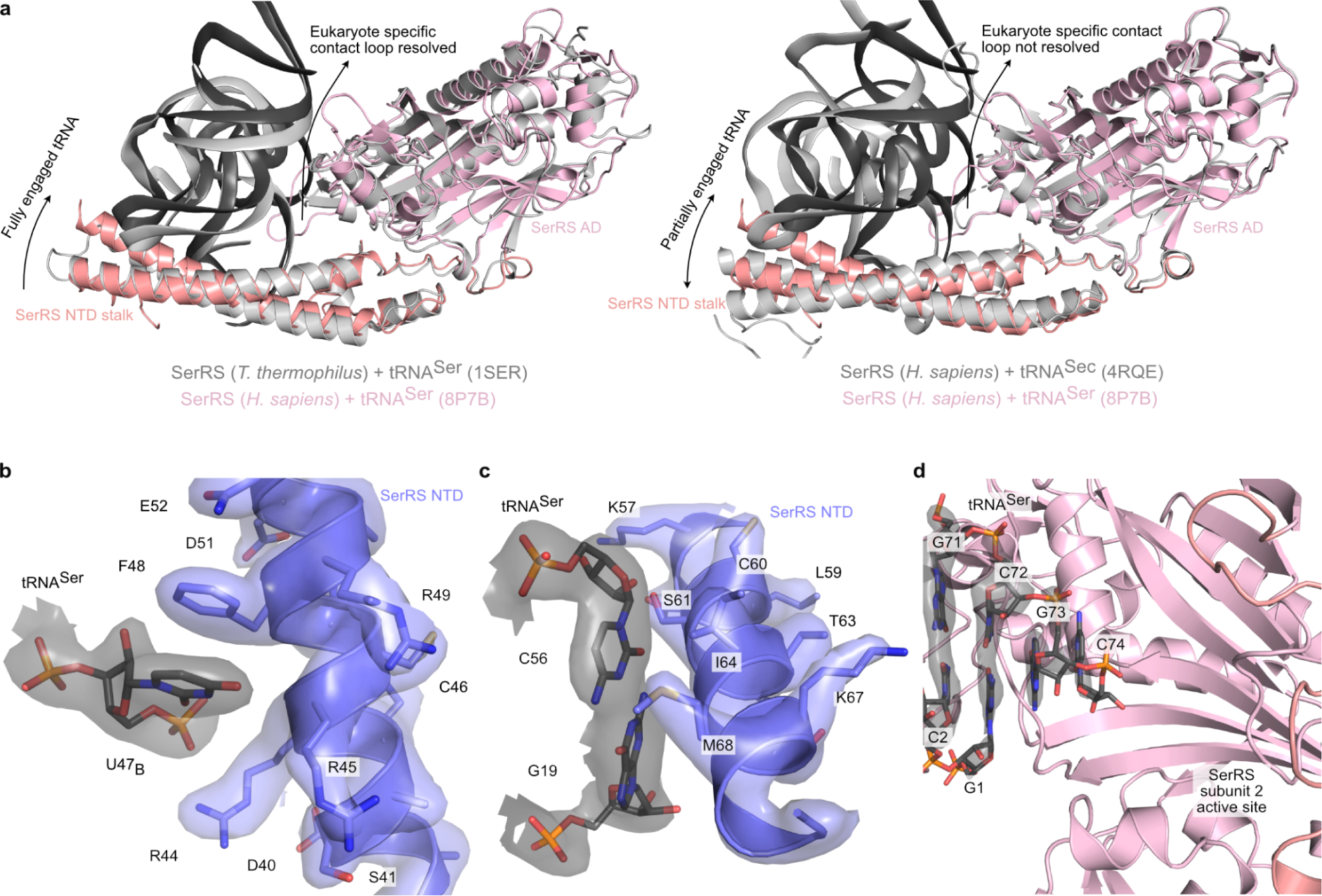
Binding of SerRS to tRNA **a** Left panel: superposition of the structure of human SerRS binding to tRNA^Ser^ (this study) with the structure of *Thermus thermophilus* SerRS bound to tRNA^Ser^ (PDB-ID: 1SER, rmsd = 1.004 Å). Left panel: superposition of the structure of human SerRS binding to tRNA^Ser^ (this study) with the structure of human SerRS binding to selenocysteine tRNA (tRNA^Sec^) (PDB-ID: 4RQE, rmsd = 0.819 Å). **b** Close-up view of the interaction between the SerRS N-terminal stalk and U47_B_ in the variable arm of tRNA^Ser^, with the cryo-EM map represented as a transparent surface. **c** Close-up view of the interaction between the SerRS N-terminal stalk and the G_19_:C_56_ base pair of tRNA^Ser^, with the cryo-EM map represented as a transparent surface. **d** Close-up view of the 3’-end of the acceptor arm of tRNA^Ser^ within the active site of SerRS, with the cryo-EM map of the tRNA acceptor arm represented as a transparent surface.

**Extended Data Figure 4:**
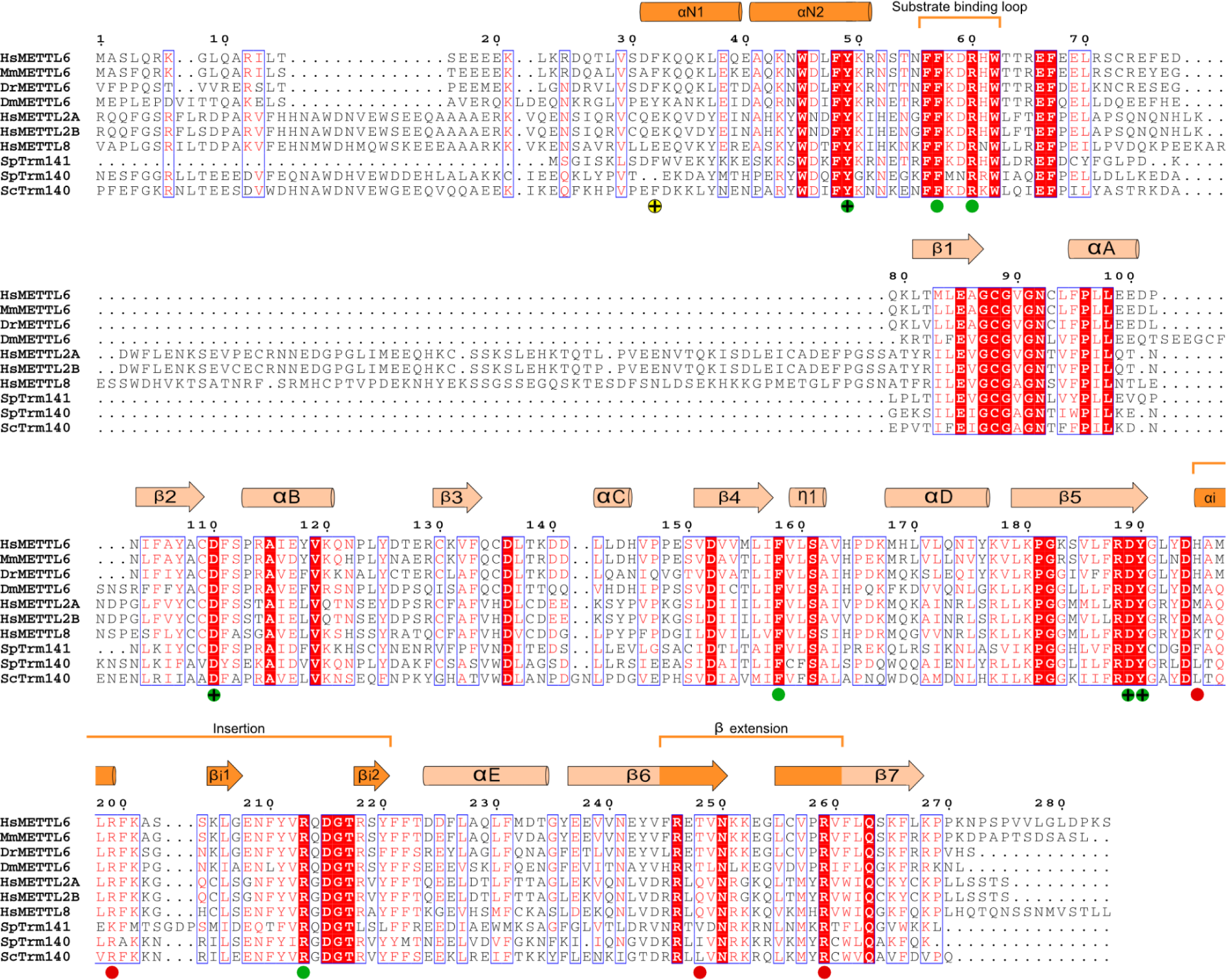
Multiple sequence alignment of m^3^C RNA methyltransferases. White letters on red background signify 100% of conservation, blue boxes indicate >70% residue similarity with the conserved residues in red. A secondary structure representation of human METTL6 from our cryo-EM model is shown at the top. Residues involved in the coordination of C_32_ are annotated with a green circle. Phenylalanine F32 in the METTL6-SerRS interface is annotated with a yellow circle. Residues involved in the coordination of i^6^A_37_ are annotated with a red circle. Single-point mutants analysed in this study are additionally annotated with a “**+**” sign.

**Extended Data Figure 5:**
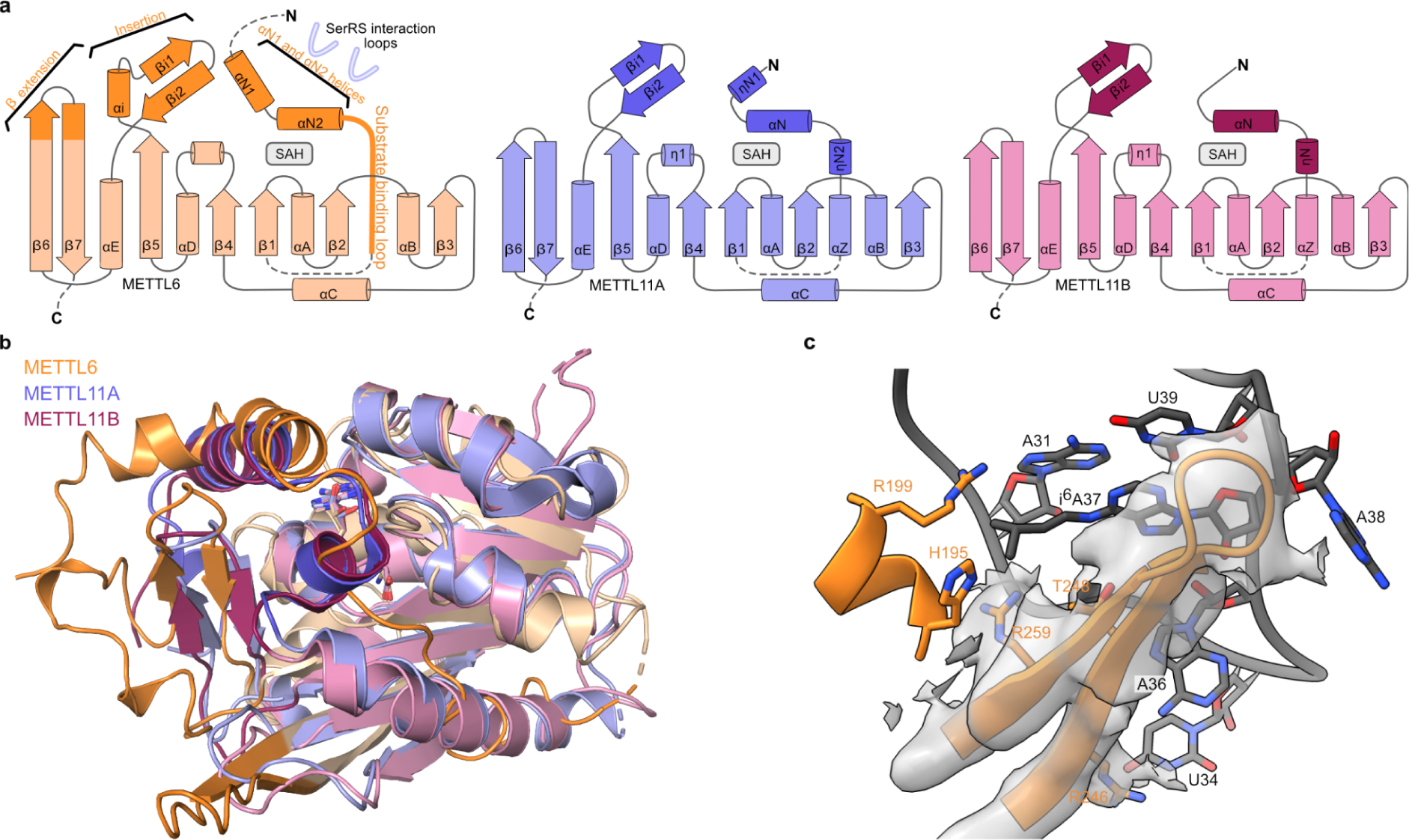
Comparison with METTL11. **a** Secondary structure diagrams of METTL6 in comparison with METTL11A and METTL11B. Dotted lines represent unmodelled parts of the experimental structures. **b** Superposition of the model of tRNA bound METTL6 with METTL11A (PDB:5UBB, rmsd = 1.137 Å) and METTL11B (PDB:6DUB) (rmsd = 1.102 Å) **c** Detailed view of the 1-extension of METTL6 with the cryo-EM map represented as a transparent surface. Low resolution in this region indicates the flexibility of the hairpin.

**Extended Data Figure 6:**
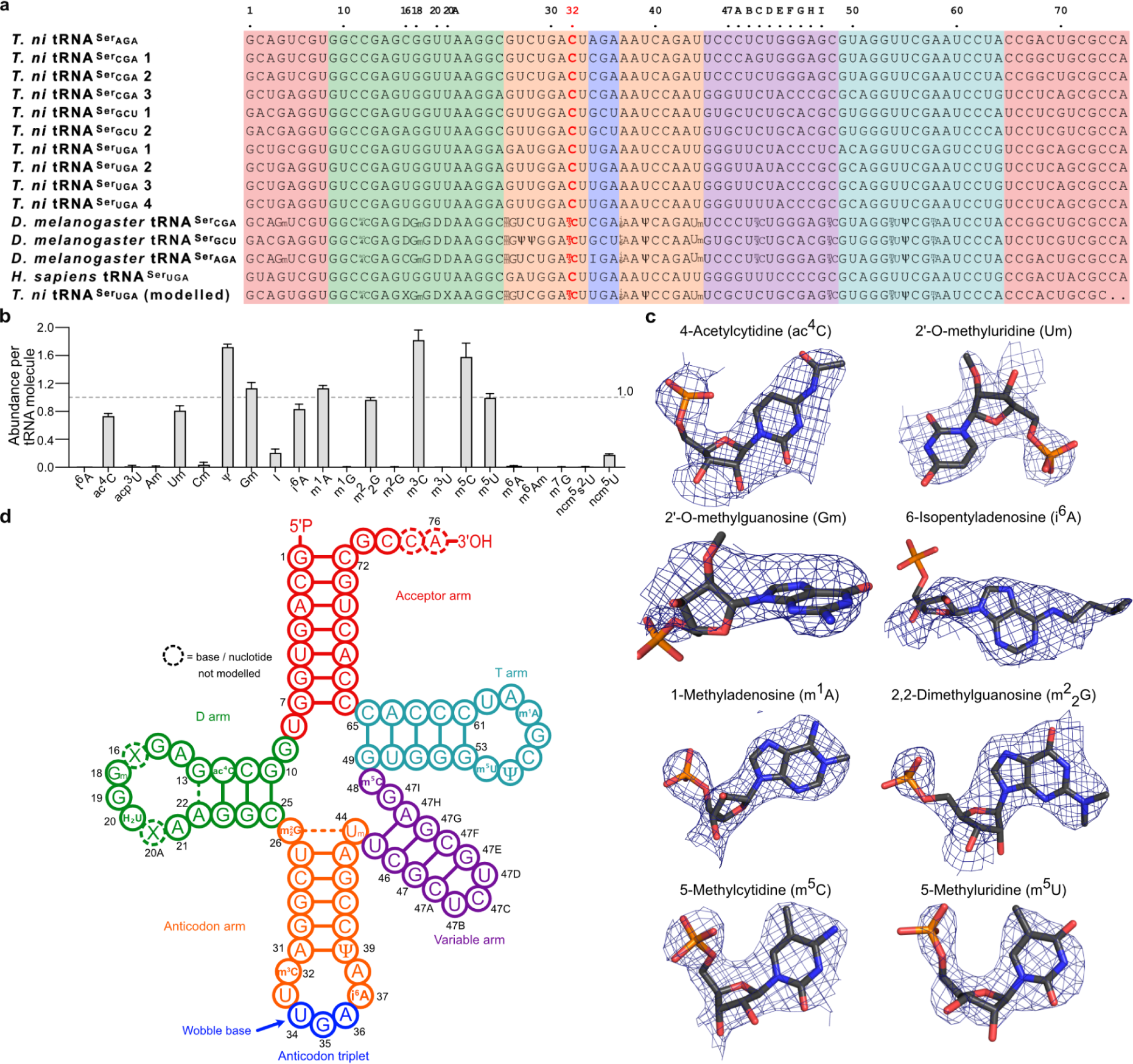
Modifications in the co-purified tRNA^Ser^ from *T. ni* insect cell expression **a** Sequence alignment of the ten tRNA^Ser^ present in expression host *T. ni*, the three tRNA^Ser^ from *D. melanogaster* with mapped modifications, *H. sapiens* tRNA^Ser^ and the consensus sequence we used for our model. The background colours represent the tRNA secondary structure features. **b** Relative abundance of modified nucleotides per tRNA molecule as determined by mass spectrometry. See also Extended Table 4. **c** Example of the cryo-EM map around modified nucleotides at positions specified in panel d. Stick model with heteroatom colour scheme. **d** Cloverleaf representation of the tRNA^Ser^ sequence in the model with colours as in panel a.

**Extended Data Figure 7:**
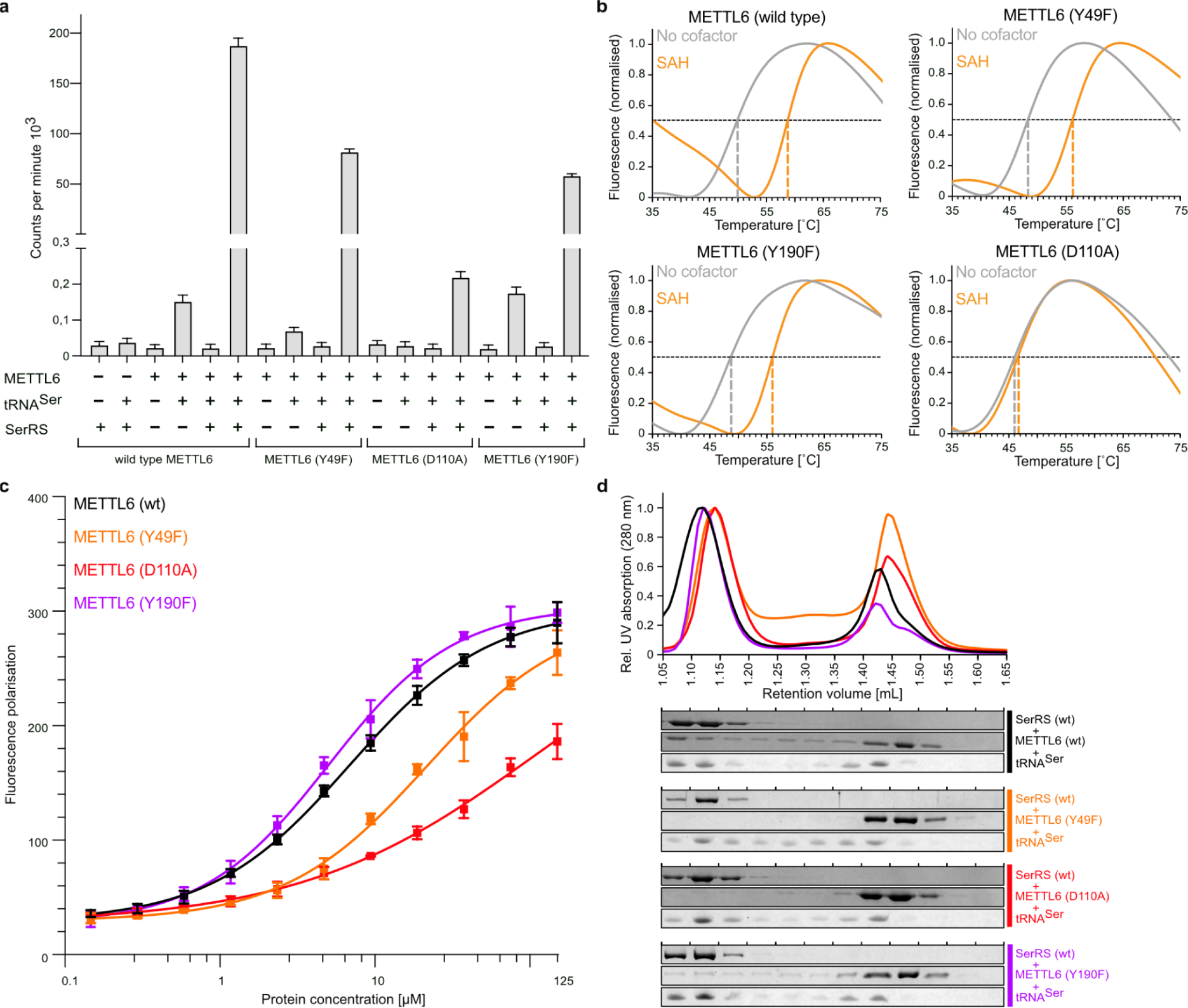
Characterization of METTL6 mutants **a** *In vitro* methylation activity measurements of the different METTL6 mutants. Data is represented as the blank-subtracted mean and standard deviation of independent replicates (n=3). **b** Thermal shift assay of METTL6 wild type and mutants, in the presence or absence of SAH. SAH binding is indicated by an increase in the melting temperature. A representative melting curve of 2 replicates is shown. **c** Interaction of the different METTL6 mutants with tRNA^Ser^ measured by fluorescence polarisation. Data is represented as the mean and standard deviation of independent replicates (n=3), and the sigmoidal fit of the curve. **d** Complex formation of SerRS mutants, METTL6 mutants and tRNA^Ser^ qualitatively analysed via analytical size-exclusion chromatography (Column: Superdex 200 3.2/300). Fractions were analysed for protein and RNA contents using SDS-PAGE and urea PAGE.

**Extended Data Figure 7:**
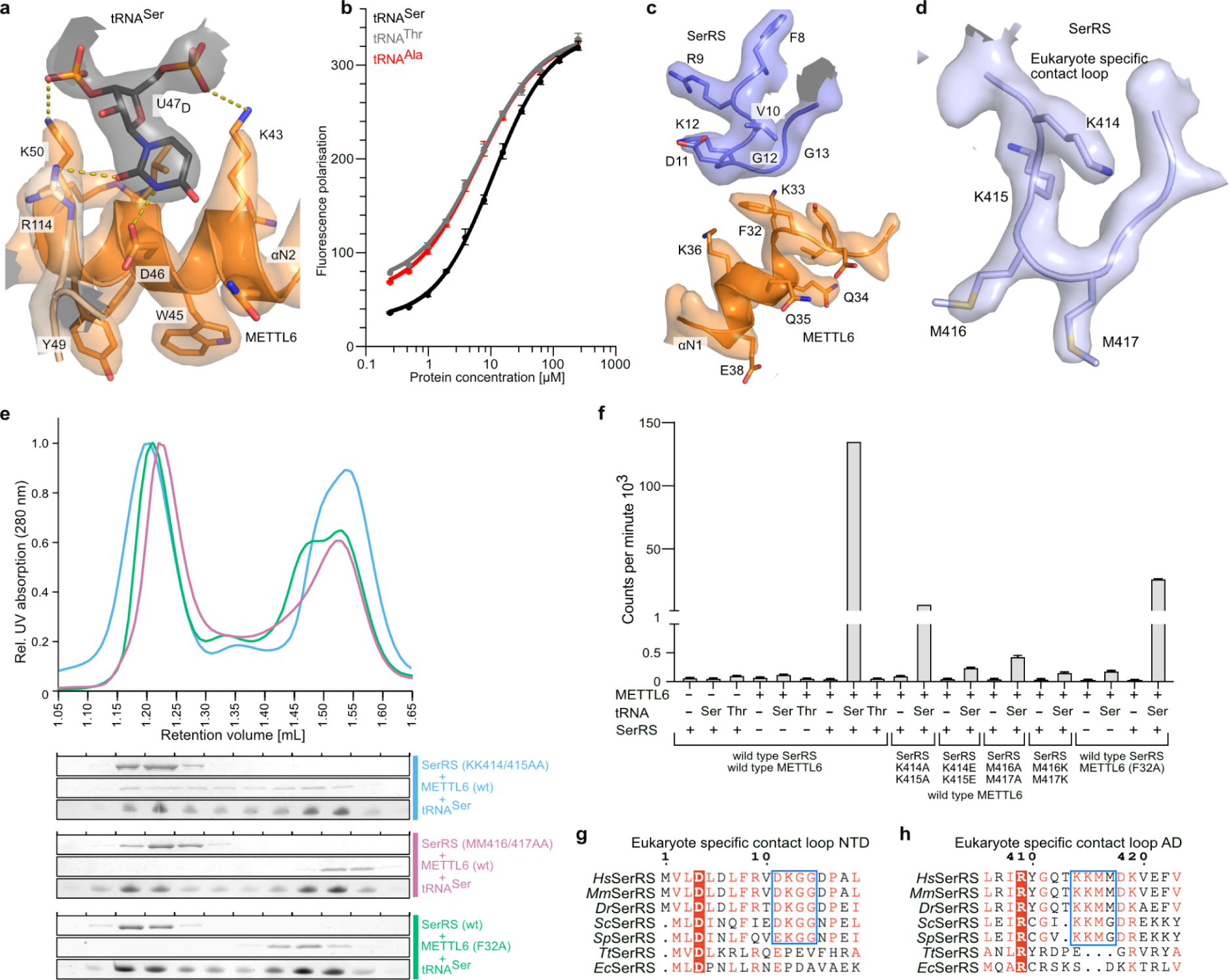
Details of METTL6 - SerRS interaction **a** Close-up view of METTL6 binding of U47_D_ with the corresponding section of the cryo-EM map. **b** Interaction of METTL6 with different tRNAs measured by fluorescence polarisation. Data is represented as the mean and standard deviation of independent replicates (n=3), and the sigmoidal fit of the curve. **c** Close-up view of the interaction between F32 of METTL6 and the N-terminal domain of SerRS with the corresponding section of the cryo-EM map. **d** Close-up view of the KKMM loop of SerRS with the corresponding section of the cryo-EM map. **e** Complex formation of SerRS mutants, METTL6 mutants and tRNA^Ser^ qualitatively analysed via analytical size-exclusion chromatography (Column: Superdex 200 3.2/300). Fractions were analysed for protein and RNA contents using SDS-PAGE and urea PAGE. **f** *In vitro* methylation activity of METTL6 and SerRS mutants on tRNA^Ser^ and tRNA^Thr^. Data is represented as the blank-subtracted mean and standard deviation of independent replicates (n=3). **g and h** Multiple sequence alignment of SerRS sequences in the METTL6 contact loops.

**Extended Table 1:** X-ray crystallographic data collection and refinement statistics.

|  | METTL6 (Full Length) – 8OWX | METTL6 (40-269) – 8OWY |
| --- | --- | --- |
| **Data collection** |  |  |
| Space group | P 4<sub>3</sub> | C 2 |
| Cell dimensions |  |  |
| *a*, *b*, *c* (Å) | 82.856, 82.856, 130.39 | 81.971, 138.129, 39.66 |
| $\alpha$, $\beta$, $\gamma$ (°) | 90, 90, 90 | 90, 115.257, 90 |
| Resolution (Å) | 41.43 - 2.601 (2.694 - 2.601) | 39.11 - 3.2 (3.315 - 3.2) |
| *R*<sub>sym</sub> or *R*<sub>merge</sub> | 0.2006 (4.8) | 0.2055 (2.196) |
| *I* / $\sigma I$ | 9.43 (0.45) | 6.97 (0.80) |
| Completeness (%) | 98.36 (83.61) | 85.31 (48.00) |
| Redundancy | 13.6 (13.3) | 6.5 (6.2) |
| **Refinement** |  |  |
| Resolution (Å) | 41.43 - 2.601 (2.694 - 2.601) | 39.11 - 3.2 (3.315 - 3.2) |
| No. reflections | 26552 (2219) | 5634 (312) |
| *R*<sub>work</sub> / *R*<sub>free</sub> | 0.2170 (0.3804)/<br>0.2503 (0.3708) | 0.2603 (0.3426)/<br>0.2990 (0.4826) |
| No. atoms | 4005 | 2673 |
| Protein | 3936 | 2623 |
| Ligand/ion | 69 | 50 |
| Water | 0 | 0 |
| *B*-factors |  |  |
| Protein | 86.58 | 51.66 |
| Ligand/ion | 77.30 | 46.26 |
| Water | N/A | N/A |
| R.m.s. deviations |  |  |
| Bond lengths (Å) | 0.012 | 0.013 |
| Bond angles (°) | 1.85 | 2.03 |

Values in parentheses are for highest-resolution shell.

**Extended Table 2:**
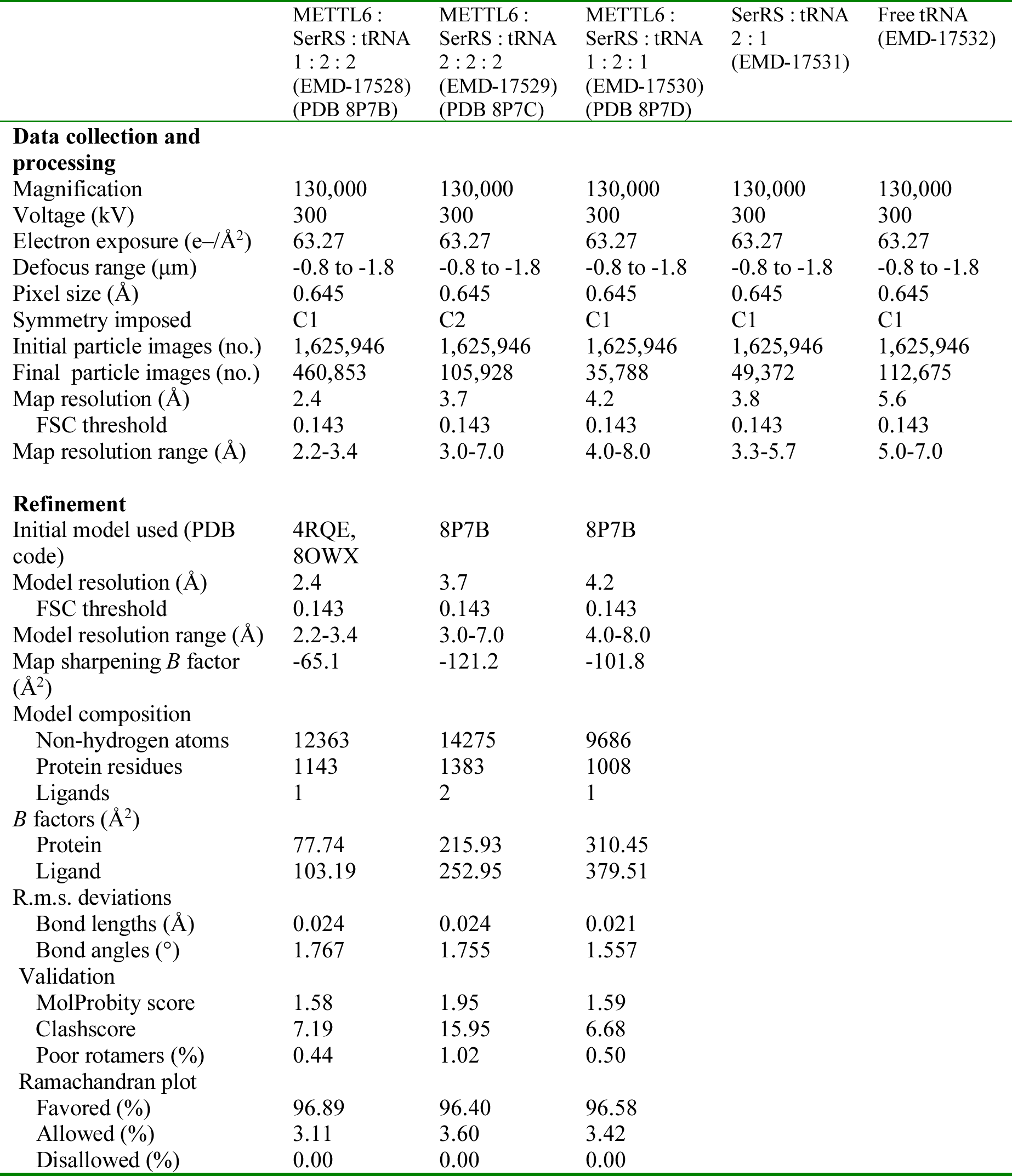
Cryo-EM data collection, refinement and validation statistics.

**Extended Table 3:** RMSD between different structures.

| Protein (PDB or AlphaFold DB accession) | RMSD (Å) against<br>METTL6 bound to tRNA<br>(8P7B) |
| --- | --- |
| METTL6 unbound full length (8OWX) | 0.654 |
| METTL6 unbound 40-269 (8OWY) | 0.637 |
| METTL6 unbound full length (7EZG) | 0.590 |
| METTL6 AlphaFold DB (Q8TCB7) | 0.611 |
| SpTrm141 AlphaFold DB (O74386) | 0.787 |
| SpTrm140 AlphaFold DB (Q9P7L6) | 0.960 |
| ScTrm140 AlphaFold DB (Q08641) | 0.985 |
| METTL2A AlphaFold DB (Q96IZ6) | 0.974 |
| METTL2B AlphaFold DB (Q6P1Q9) | 0.992 |
| METTL8 AlphaFold DB (B3KW44) | 0.916 |

**Extended Table 4:** MS/MS parameters for quantification of modified nucleoside.

| Compound | Precursor Ion | Product Ion | Rt (min) | ΔRt (min) | Fragmentor (V) | Collision Energy (eV) |
| --- | --- | --- | --- | --- | --- | --- |
| A | 268,1 | 136 | 5,2 | 1 | 200 | 20 |
| A SILIS | 283 | 146 | 5,2 | 1 | 200 | 20 |
| ac4C | 286,1 | 154 | 5 | 1 | 85 | 9 |
| ac4C SILIS | 300 | 163 | 5 | 1 | 85 | 9 |
| acp3U | 346,1 | 214,1 | 2,3 | 1 | 95 | 15 |
| Am | 282,1 | 136 | 6 | 1 | 130 | 17 |
| Am SILIS | 298 | 146 | 6 | 1 | 130 | 17 |
| C | 244,1 | 112 | 2,1 | 1 | 200 | 20 |
| C SILIS | 256 | 119 | 2,1 | 1 | 200 | 20 |
| Cm | 258,1 | 112 | 4,1 | 1 | 180 | 9 |
| Cm SILIS | 271 | 119 | 4,1 | 1 | 180 | 9 |
| G | 284,1 | 152 | 4,3 | 1 | 200 | 20 |
| G SILIS | 299 | 162 | 4,3 | 1 | 200 | 20 |
| Gm | 298,1 | 152 | 5 | 1 | 100 | 9 |
| Gm SILIS | 314 | 162 | 5 | 1 | 100 | 9 |
| I | 269,1 | 137 | 4,1 | 1 | 100 | 10 |
| I SILIS | 283 | 146 | 4,1 | 1 | 100 | 10 |
| i6A | 336,3 | 204,1 | 8 | 1 | 140 | 17 |
| i6A SILIS | 356 | 219 | 8 | 1 | 140 | 17 |
| m1A | 282,1 | 150 | 2,5 | 2 | 150 | 25 |
| m1A SILIS | 298 | 161 | 2,5 | 2 | 150 | 25 |
| m1G | 298,1 | 166 | 4,9 | 1 | 105 | 13 |
| m1G SILIS | 314 | 177 | 4,9 | 1 | 105 | 13 |
| m1I | 283,1 | 151 | 4,8 | 1 | 80 | 12 |
| m1I SILIS | 298 | 161 | 4,8 | 1 | 80 | 12 |
| m1Y | 259 | 223 | 3,1 | 1 | 85 | 5 |
| m22G | 312,1 | 180 | 5,7 | 1 | 105 | 13 |
| m22G SILIS | 329 | 192 | 5,7 | 1 | 105 | 13 |
| m2G | 298,1 | 166 | 5,1 | 1 | 95 | 17 |
| m2G SILIS | 314 | 177 | 5,1 | 1 | 95 | 17 |
| m3C | 258,1 | 126 | 2,3 | 1,5 | 88 | 14 |
| m3C SILIS | 271 | 134 | 2,3 | 1,5 | 88 | 14 |
| m3U | 259,1 | 127 | 4,8 | 0,6 | 75 | 9 |
| m5C | 258,1 | 126 | 3,8 | 1 | 185 | 13 |
| m5C SILIS | 271 | 134 | 3,8 | 1 | 185 | 13 |
| m5U | 259,1 | 127 | 4,4 | 1 | 95 | 9 |
| m5U SILIS | 271 | 134 | 4,4 | 1 | 95 | 9 |
| m66A | 296 | 164 | 7,1 | 1 | 130 | 21 |
| m66A SILIS | 313 | 176 | 7,1 | 1 | 130 | 21 |
| m6A | 282,1 | 150 | 6,5 | 1 | 125 | 17 |
| m6A SILIS | 298 | 161 | 6,5 | 1 | 125 | 17 |
| m7G | 298,1 | 166 | 3,6 | 1 | 100 | 13 |
| m7G SILIS | 314 | 177 | 3,6 | 1 | 100 | 13 |
| mcm5s2U | 333,1 | 201 | 6,2 | 1 | 92 | 8 |
| mcm5s2U SILIS | 347,1 | 210 | 6,2 | 1 | 92 | 8 |
| ncm5s2U | 318,1 | 186 | 4,2 | 1 | 95 | 7 |
| mcm5U | 317,1 | 185,1 | 5 | 1 | 95 | 5 |
| mcm5U SILIS | 331 | 194 | 5 | 1 | 95 | 5 |
| ncm5U | 302 | 170 | 2,5 | 1 | 85 | 8 |
| ncm5U SILIS | 316 | 179 | 2,5 | 1 | 85 | 8 |
| Q | 410,2 | 295,1 | 4,3 | 1 | 115 | 12 |
| t6A | 413,1 | 281,1 | 5,8 | 1 | 130 | 9 |
| t6A SILIS | 434 | 297 | 5,8 | 1 | 130 | 9 |
| U | 245,1 | 113 | 3 | 1 | 95 | 5 |
| U SILIS | 256 | 119 | 3 | 1 | 95 | 5 |
| Um | 259,2 | 113 | 4,6 | 1 | 96 | 8 |
| Um SILIS | 271,1 | 119 | 4,6 | 1 | 96 | 8 |
| Y | 245,1 | 209 | 1,7 | 1 | 90 | 5 |
| Y SILIS | 256 | 220 | 1,7 | 1 | 90 | 5 |
| ManQ | 572,3 | 295,5 | 3,9 | 1 | 120 | 20 |
| GalQ | 572,3 | 295,5 | 4,1 | 1 | 115 | 20 |

